# Brain network dynamics predict moments of surprise across contexts

**DOI:** 10.1101/2023.12.01.569271

**Authors:** Ziwei Zhang, Monica D. Rosenberg

## Abstract

We experience surprise when reality conflicts with our expectations. When we encounter such expectation violations in psychological tasks and daily life, are we experiencing completely different forms of surprise? Or is surprise a fundamental psychological process with shared neural bases across contexts? To address this question, we identified a brain network model, the surprise edge-fluctuation-based predictive model (EFPM), whose regional interaction dynamics measured with functional magnetic resonance imaging (fMRI) predicted surprise in an adaptive learning task. The same model generalized to predict surprise as a separate group of individuals watched suspenseful basketball games and as a third group watched videos violating psychological expectations. The surprise EFPM also uniquely predicts surprise, capturing expectation violations better than models built from other brain networks, fMRI measures, and behavioral metrics. These results suggest that shared neurocognitive processes underlie surprise across contexts and that distinct experiences can be translated into the common space of brain dynamics.

---

Picture this scenario. You walk into a restaurant anticipating a regular get-together with friends and are shocked when the crowd shouts out “happy birthday!”, revealing that it is a planned party. In a similar vein, imagine gasping at an unexpected move in a sporting event or a plot twist in a movie. Surprise is a fundamental human experience, but how much do these surprising moments in our life have in common? Despite being in completely different situations, does our brain process unexpected changes in those moments similarly?

Formally, surprise scales with the difference between our prediction about the world and the actual outcome we observe: the bigger the difference, the bigger the surprise. In reality, we may be more surprised by prediction errors with practical significance. Previous work suggests that outcomes substantial enough to shift observers’ internal model of the current situation result in belief-inconsistent surprise (Baldi & Itti, 2010). The more a piece of evidence violates our current belief, the higher the belief-inconsistent surprise. For example, your belief about the purpose of the get-together with friends was violated after the surprising moment.

Psychologists quantify belief-inconsistent surprise as the degree to which an observation has shifted belief about the environment. Tasks used to measure belief-inconsistent surprise usually involve participants making adaptive predictions in a changing environment and have revealed a set of brain structures that encode information about surprise. Univariate functional magnetic resonance imaging (fMRI) activation in the default mode network (e.g., posterior cingulate cortex), frontoparietal network (e.g., posterior parietal cortex [McGuire et al., 2014], right superior temporal sulcus and frontal eye fields [Meyniel & Dehaene, 2017]), and limbic system (e.g., dorsal cingulate cortex [O’Reilly et al., 2013]) have been shown to track surprise. Beyond fMRI activation, Kao et al. (2020) related belief-shifting changes in an adaptive learning task (McGuire et al. 2014) to functional connectivity dynamics between frontoparietal and other large-scale brain networks.

Functional connectivity dynamics have typically been measured with approaches that calculate statistical dependence between two regions’ BOLD signal time series over windows of time on the order of tens of seconds (Allen et al., 2014; Lurie et al., 2020; Preti et al., 2017). Recent advances in network neuroscience have made it possible to investigate higher-frequency interaction dynamics by computing two regions’ moment-by-moment co-deflections, known as their co-fluctuation or edge time series (Faskowitz et al. 2020; Zamani Esfahlani et al., 2020). Co-fluctuation patterns calculated from the whole brain allow us to capture brain-wide interactions and do so at the level of single frames. This is a parameter-free method to capture dynamic functional interactions that does not involve calculating the correlation between regions within sliding windows of a selected size (Betzel et al., 2023; Tanner et al., 2023). These advances are especially useful for capturing processes such as surprise that tend to be transient (e.g., the surprising “happy birthday!” might elicit a strong but instantaneous response).

Previous work has found associations between surprise and brain network dynamics, but an open question is whether surprise in very different situations is subserved by similar neurocognitive processes. This is difficult to assess with behavioral measures alone because in some paradigms surprise is measured explicitly (e.g., via button presses) whereas in others it is hidden (e.g., during passive viewing). Although other measures such as pupil size (Kloosterman et al., 2015; Lavin et al., 2014; Liao et al., 2018; Preuschoff et al., 2011) and facial expression (Chang et al., 2021) track surprise in some contexts, they may be confounded by low-level visual properties or less sensitive to surprise in others.

Characterizing brain dynamics allows us to discover commonalities between belief-inconsistent surprise in distinct contexts. To this end, we propose an edge-fluctuation-based predictive model (EFPM) trained to identify functional interactions predicting moment-to-moment changes in belief-inconsistent surprise. This model, the surprise EFPM, predicts surprise across datasets and is uniquely generalizable when compared with models based on other functional brain networks and BOLD activation alone as well as models built to predict other behavioral measures. Theoretically, this result reveals shared neurocognitive processes between belief-inconsistent surprise in task and naturalistic contexts. Practically, the approach offers a new way to predict cognitive dynamics from brain dynamics and translate different experiences into the shared space of the brain.

## Results

### An edge-fluctuation-based predictive model (EFPM) predicts belief-inconsistent surprise in a controlled learning task

Not all new observations are equally informative and meaningful. Some observations differ substantially from what was predicted and thus induce belief-inconsistent surprise—that is, surprise that changes our belief about the world (Itti & Baldi, 2009). Belief-inconsistent surprise drives learning (Sutton & Barto, 2018) and demarcates subjective experiences (Antony et al., 2021; Kumar et al., 2022).

Here we aim to identify neural predictors of belief-inconsistent surprise. To do so, we first reanalyzed an openly available MRI dataset (*n*=32) in which participants performed an adaptive learning task during functional imaging (McGuire et al., 2014; Kao et al., 2020). Participants’ goal was to predict the location of an upcoming object (a dropped bag). The location of the bag was drawn from a generative distribution defined by a mean (a “helicopter” position) and standard deviation hidden from participants (**Fig. 1**). On each trial, participants predicted the location of the bag by moving a “bucket” below where they thought it would fall. The bag then dropped and exploded into coins. Coins that fell into the bucket, indicating the precision of the prediction, and were added to a cumulative score. After each bag drop, a red bar indicated the gap between the bag’s location and the participant’s earlier prediction. Importantly, the mean of the hidden generative distribution remained unchanged most of the time. However, the probability of the mean being randomly redrawn from a uniform distribution was 0.125 on each trial. A change trial is a sudden, unpredictable, and surprising event in the environment.

**Figure 1.**
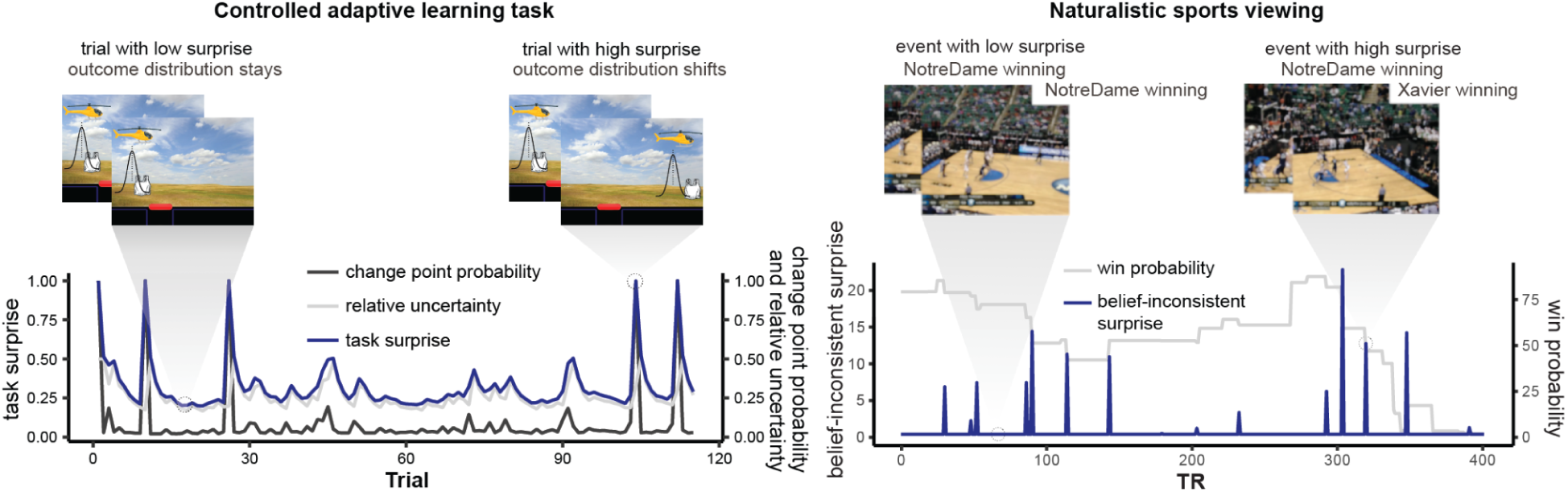
Task overview. **Left**: The belief-inconsistent surprise time course in the controlled adaptive learning task is visualized using data shared by McGuire et al., 2014. Examples of trials with high surprise, when a change point has likely occurred, and low surprise, when a change point was not likely, are shown. Note that participants were not shown with the helicopter during the main task but were aware of its existence from practice. Task figures were recreated here for visualization purposes. **Right**: Belief-inconsistent surprise time course in the naturalistic sports viewing paradigm visualized from data associated with Antony et al., 2021. Surprise occurs when a predicted winner becomes more likely to lose and a predicted loser becomes more likely to win. Video screenshots are blurred for copyright reasons.

McGuire et al. (2014) defined a normative model that describes the optimal way of performing this task. To update their prediction on the next trial, an ideal learner would take into account two factors: 1) the probability that the generative mean has shifted given the current outcome and 2) the uncertainty about whether there is a true shift in the generative mean or simply noise around the previous generative mean. To quantify these factors, they developed two measures, change-point probability (CPP) and relative uncertainty (RU). This learning process can be described by the delta learning rule where belief about the helicopter location is updated using the difference between the observed outcome and current prediction, multiplied by a learning rate (model described in detail in Nassar et al., 2012, 2010). However, an ideal learner should know that not all outcomes warrant updating predictions to the same extent. Rather, moments of high surprise are more likely to signal a change in the underlying environment (e.g., the hidden distribution) and should thus lead to more updating. Therefore, the learning rate, which signals how much an ideal learner should update their belief about the hidden distribution after seeing an outcome, was formalized as CPP+RU*(1-CPP). We use this compound measure from change-point probability and relative uncertainty, calculated and shared by McGuire et al. (2014) and Kao et al. (2020), as our measure of belief-inconsistent surprise. Participants’ belief is equivalent to the mean of the generative distribution and belief-inconsistent surprise is thus higher when there is a mean shift.

How can we capture the neural predictors of surprise in a highly dynamic environment? Work using controlled tasks and naturalistic paradigms has demonstrated that fluctuations in other cognitive processes, such as sustained attention (Rosenberg et al., 2020; Jones et al., 2023), working memory (Shine et al., 2016), decision making (Taghia et al., 2018), learning (Bassett et al., 2011), and narrative engagement (Song et al., 2021), are reflected in the dynamics of functional brain connections, or edges. Edge dynamics have traditionally been measured by correlating two regions’ activity time courses in time windows. Windowed approaches, however, may not be fine-grained enough to capture surprise dynamics, which can change rapidly. Here we track the network predictors of surprise using a new approach developed by Faskowitz et al. (2020) and Zamani Esfahlani et al. (2020) that decomposes edge strength into moment-by-moment co-fluctuation values. Co-fluctuation between every pair of regions in the brain is computed by taking the product of their *z*-scored time series, resulting in an edge time series (Faskowitz et al. 2020; Zamani Esfahlani et al., 2020). Here we ask if edge time series predict momentary changes in belief-inconsistent surprise. To do so, we implemented a new method, edge-fluctuation-based predictive modeling (EFPM).

Using leave-one-subject-out cross-validation, we identified edges whose strength varied across trials with belief-inconsistent surprise (**Supp Fig. 1**). In each training fold, we selected 31 (*n*-1) participants and calculated the partial Spearman correlation (*rho*) between their edge time series and their belief-inconsistent surprise time course, controlling for head motion (framewise displacement, FD; see *Methods*) at every trial. We Fisher *z*-transformed *rho* values, conducted a one-sample *t* test comparing these 31 *rho* values against zero, and selected edges significantly positively and negatively correlated with belief-inconsistent surprise across the group at *p*<0.05. In the held-out individual, we separately averaged the time series of these edges positively and negatively correlated with surprise. We calculated the partial *rho* between these time courses and the held-out subject’s belief-inconsistent surprise time course, controlling for framewise displacement at every trial. This process was repeated so that each individual was held out once. To assess the significance of the correlation between the edge and surprise time series we conducted a non-parametric test comparing the observed mean partial *rho* values to the means of null partial *rho* distributions obtained via permutation testing (see *Methods*).

EFPM successfully predicted task surprise in held-out individuals (**Fig. 2A**). Edges positively correlated with surprise were stronger on trials with more unexpected outcomes (mean within-subject partial *rho*=0.09; two-tailed *p*=1/1001) whereas edges negatively correlated with surprise showed the opposite pattern (mean within-subject partial *rho*=-0.10; two-tailed *p*=1/1001). Predictions remained significant when permutation testing was performed by randomly selecting same-size edge sets for prediction (*p* values=1/1001) rather than phase-randomizing the mean time series of edges positively and negatively correlated with surprise. Thus, moment-to-moment changes in edge strength predict moment-to-moment changes in belief-inconsistent surprise in novel individuals.

**Figure 2.**
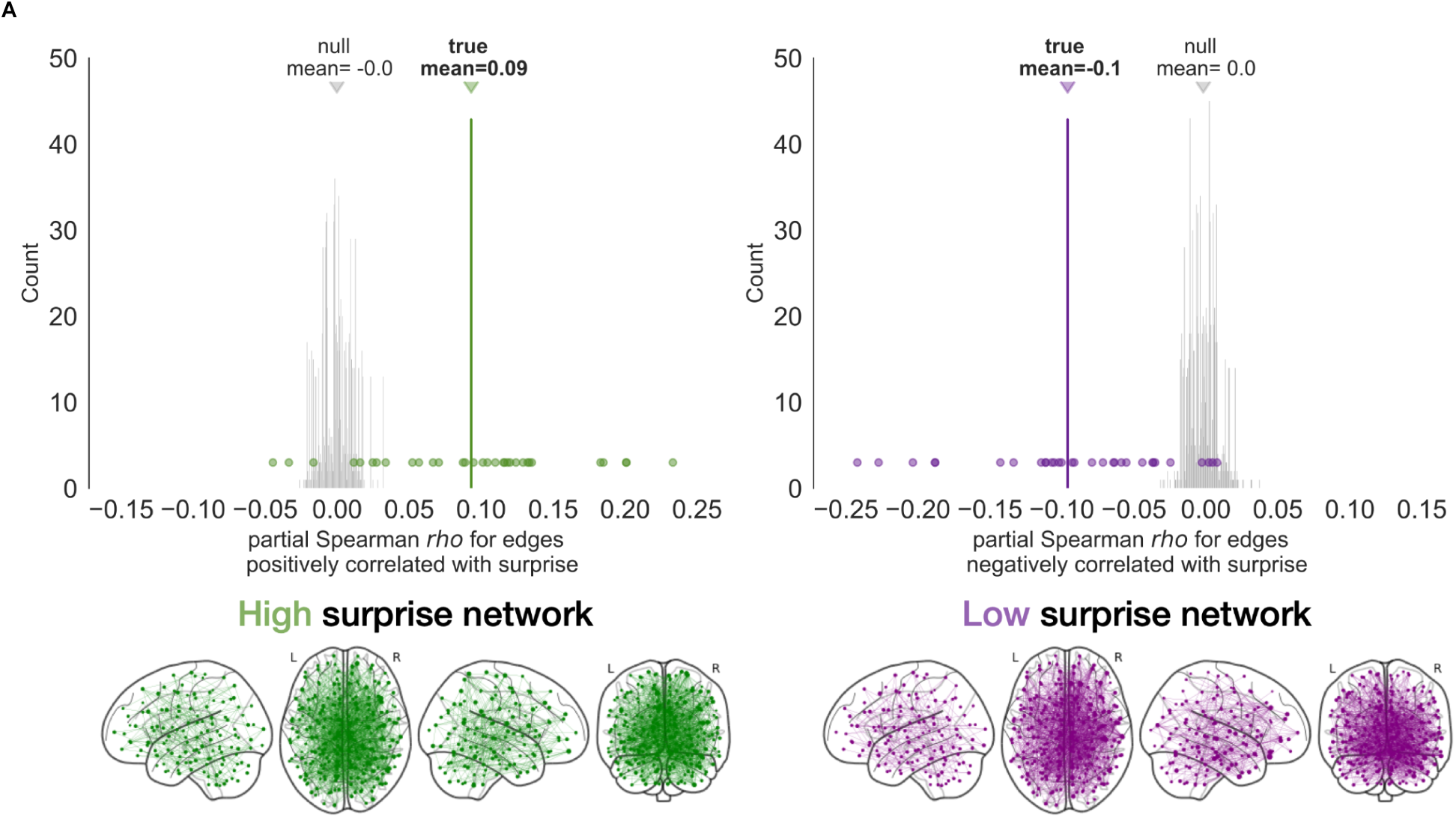
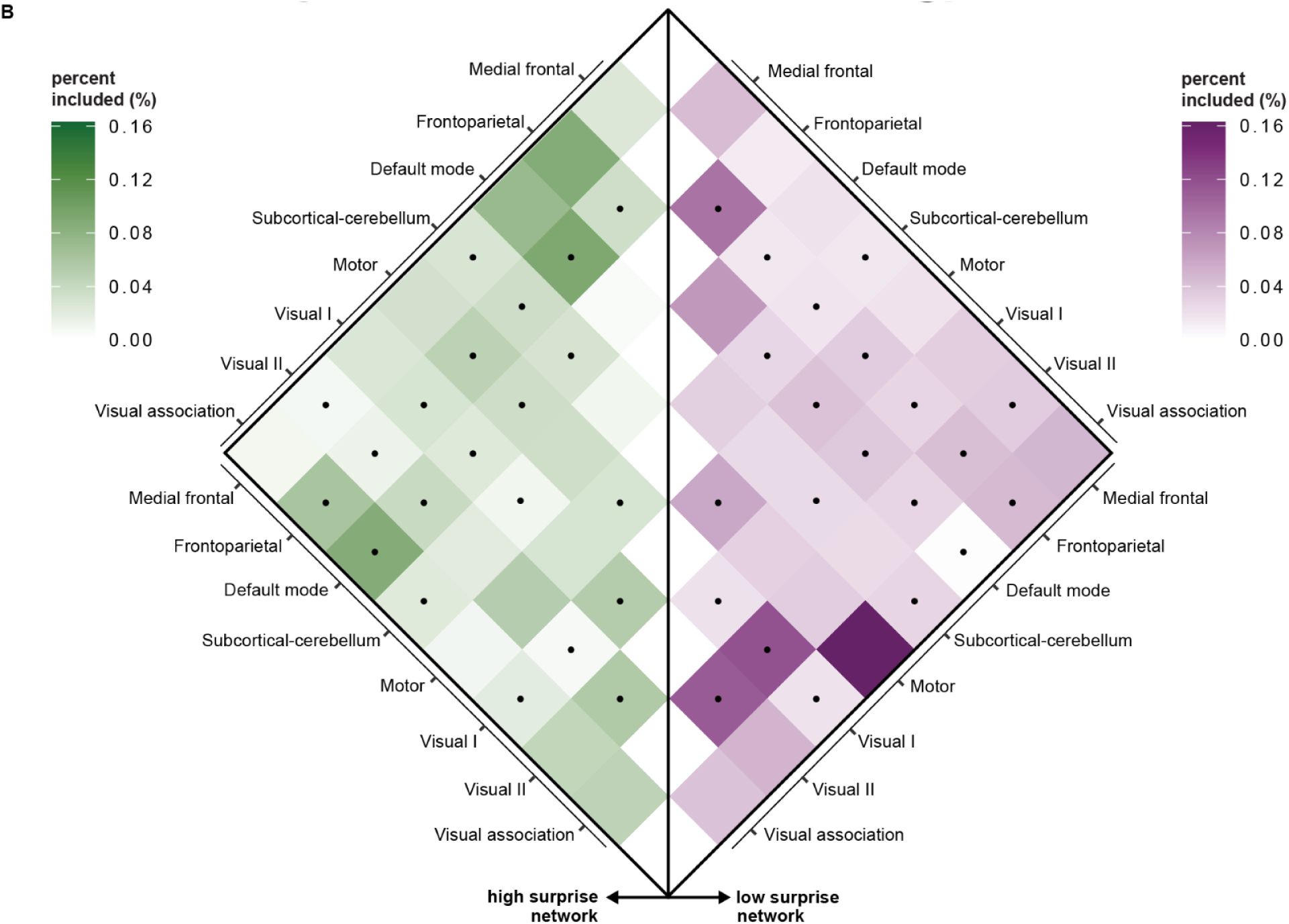
Surprise EFPM. Results from training the edge-fluctuation-based predictive model of surprise. **A**, The observed partial Spearman *rho* value for the high (left, where each green dot represents a participant) and low (right, where each purple dot represents a participant) surprise network was plotted with the null distribution (gray distribution) obtained from repeating the internal cross-validation 1000 times with phase-randomized time series of the strength of the network positively and negatively correlated with surprise, respectively, and then averaged across participants (**top**). Edges that appeared in every fold of the cross-validation process were selected into the surprise EFPM and visualized (**bottom**). **B**, Surprise EFPM anatomy. Edges in both high and low surprise networks span predefined functional brain networks. “Percent included” indicates the percentage of edges within each functional network or between each pair of networks that belong to the high or low surprise networks, respectively. Dots show the importance of each network in cross-dataset generalization, assessed with the computational lesioning analysis.

We then defined the *high* surprise and *low* surprise networks separately as edges that appeared in all 32 leave-one-out cross-validation folds. This internal cross-validation process resulted in 1115 edges in the *high* and 1216 edges in the *low* surprise network (**Fig. 2A; Supp Fig. 2**). We examined the network anatomy by assigning edges in the surprise EFPM to canonical resting-state networks (Finn et al., 2015; **Supp Fig. 3** and **Supp Table 1**). After controlling for network size, the canonical networks most represented in the surprise model were the medial frontal (MF), frontoparietal (FP), and default mode (DMN) networks for the high surprise network and motor (MT), visual I (VI), visual II (VII) networks for the low surprise network (**Fig. 2B**).

These findings complement previous work demonstrating that univariate activations in key regions in the parietal lobe were associated with surprise during this learning task (McGuire et al., 2014). Previous time-averaged functional connectivity analyses also revealed that edges within the frontoparietal network and between the frontoparietal and other distributed networks were associated with surprise (Kao et al., 2020).

### The surprise EFPM generalizes to predict belief-inconsistent surprise during naturalistic sports viewing

Surprise EFPM network dynamics predict how an ideal learner should update predictions in the helicopter task. To test whether the surprise EFPM tracks belief-inconsistent surprise *in general* or optimal helicopter task performance *in particular*, we asked whether the *same* high and low surprise networks predict a completely different measure of surprise in a completely different context.

As we watch suspenseful sports games, we constantly generate predictions about the outcome. Basketball games, for example, are full of surprising moments when something unexpected happens on the court. If you are watching the home team dominate the game, seeing them score another free throw may not be very surprising. However, if they suddenly gain possession of the ball and score when they are losing badly, you may be more surprised.

To formally estimate surprise during U.S. National Collegiate Athletic Association (NCAA) basketball game watching, Antony et al. (2021) built a model predicting the expected probability of the home team winning each game that participants watched in the scanner. Their win-probability model was based on four factors: the score difference between the teams (designed to be positive when the ‘home’ team is leading and negative when they are behind), the remaining time, ball possession, and a team strength adjustment. Surprise was operationalized as the absolute value of the change in this win-probability time course. Belief-inconsistent surprise was presumed to occur when a change in a team’s win probability was not in line with the belief about their chance of winning. Belief-inconsistent surprise was higher when the winning team was less likely to win at each given moment and was zero otherwise. For example, consider a situation in which Notre Dame has a 90% chance of winning and Xavier scores, resulting in a 85% win probability for Notre Dame. The belief-inconsistent surprise at this moment would be 0.05. On the other hand, if Notre Dame had scored and increased their win probability to 95%, belief-inconsistent surprise would be zero since the predominant belief about Notre Dame’s chance of winning did not change. The belief-inconsistent surprise here thus conceptually resembles that in the adaptive learning task but is quantified in a naturalistic context.

We tested whether the surprise EFPM identified in the adaptive learning task generalized to predict belief-inconsistent surprise during naturalistic basketball viewing. To do so, we calculated all naturalistic-viewing participants’ edge time series and separately averaged the time series in the high and low surprise networks (see *Methods*; **Supp Fig. 1**). We calculated a surprise EFPM summary score by subtracting the average time course of the low surprise network from that of the high surprise network. This resulted in a moment-to-moment time course of surprise EFPM strength for each participant and game.

We ran a linear mixed-effects model using the surprise EFPM strength time course to predict belief-inconsistent surprise in the videos. We included other regressors as predictors (the amount of game remaining in seconds of game time at the start of the possession; which team was in possession of the ball; the auditory envelope; global video luminance; global video motion; commentator prosody; court position of the ball [left or right]; framewise displacement) to control for the variables that might co-vary with surprise. Significance was assessed with permutation testing (see *Methods*).

The surprise EFPM predicted belief-inconsistent surprise in the NCAA basketball videos (*ß*=0.037*, t*(65136.852)=3.947, *p*=0.047; for full model results, see **Supp Tables 2 and 3**), accounting for the nuisance regressors. In other words, the higher the co-fluctuation strength in the surprise EFPM, the more belief-inconsistent surprise there is in the video. The surprise EFPM, therefore, predicts belief-inconsistent surprise in general rather than helicopter task performance in particular.

We performed a computational lesioning analysis to further examine the anatomy of the surprise EFPM. For each predefined network, we first excluded all edges that involved at least one node in this network from both the high and low surprise networks, and then calculated the score of the remaining edges in the same way we calculated the network strength for the intact surprise EFPM. If a predefined network plays an important role in predicting belief-inconsistent surprise, we would expect the lesioned network to fail to generalize to the basketball-viewing dataset. Results revealed that the lesioning of edges in a range of networks (e.g., DMN–frontoparietal) led to a failure of surprise EFPM generalization and included all connections with frontoparietal network nodes (**Fig. 2B**). These results suggest that dynamics in these networks contain context-general rather than context-specific signals of surprise.

Finally, to further assess the generalizability of models built with EFPM, we flipped the training and testing set: we trained an EFPM on naturalistic belief-inconsistent surprise following the same procedure above and tested whether the model generalized to predict belief-inconsistent surprise in the learning task. An EFPM built from the flipped procedure also generalized, where the strength of edges that predicted belief-inconsistent surprise in the video also predicted surprise in the adaptive learning task (**Supp Fig. 4**). Demonstrating specificity, the strength of these edges was not related to participants’ behavioral prediction of the bag location (*ß*=-0.234*, t*(14882.818)=-0.369, *p*=0.812) or the reward they received on each trial (quantified by the number of coins participants caught in their prediction basket: *ß*=-0.399*, t*(14931.395)=-1.020, *p*=0.325; quantified by the score gained from the coins caught: *ß*=-0.551*, t*(14957.082)=-0.957, *p*=0.338). Both the high and low surprise networks identified from the two training processes significantly overlapped, further demonstrating consistency in neural systems encoding information about belief-inconsistent surprise (**Fig. 3**; *p*=0.001 for high surprise network, *p*=0.002 for low surprise network). Occipital–parietal and prefrontal-limbic edges consistently predict high surprise and intra-occipital, parietal–motor, and parietal–insular edges consistently predict low surprise.

**Figure 3.**
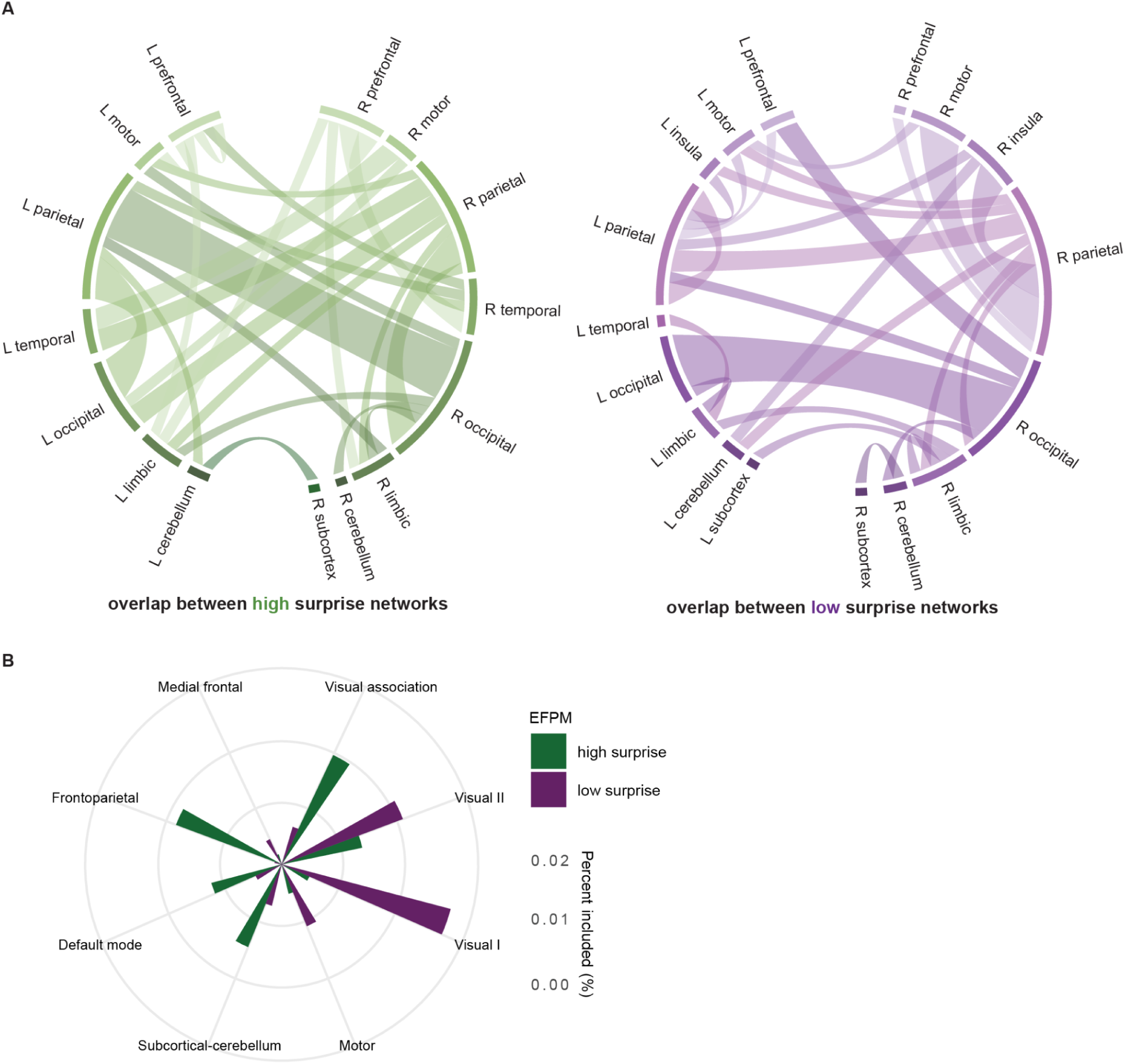
Edges that predict high or low belief-inconsistent surprise in both the task and video datasets. **A**, Anatomy of overlapping edges of each network by lobe. Thicker lines indicate that more edges in those connected networks overlap across the EFPMs identified from both datasets. **B**, Summary of the importance of each canonical resting-state network in the overlapping edges. Longer bars indicate that a higher percent of edges in that network were present in the overlapping edges.

### The surprise EFPM outperforms models built from sliding-window functional connectivity

To what extent do belief-inconsistent surprise predictions rely on the use of edge co-fluctuation, a relatively fine-grained temporal measure? To address this question, we built models based on dynamic functional connectivity calculated with sliding windows, a more traditional approach to measuring brain network dynamics (Brandman et al., 2021; Hindriks et al., 2016; Hutchison et al., 2013; Meer et al., 2020; Simony et al., 2016; Song et al., 2021). We implemented the same leave-one-subject-out model training procedures we used to obtain the surprise EFPM on dynamic functional connectivity data in the adaptive learning task calculated with three different window sizes (11, 21, 31 TRs, corresponding to 27.5, 52.5, and 77.5 seconds, respectively).

None of these sliding-window-based models significantly predicted surprise in left-out participants (**Supp Table 4**). Furthermore, no sliding-window connectivity models based on edges that overlapped across all folds of leave-one-subject-out cross-validation in the learning task dataset generalized to predict surprise in the basketball dataset (**Supp Table 4**). These results suggest that, compared with sliding-window dynamic connectivity, moment-to-moment edge fluctuations uniquely capture surprise.

### Models built from BOLD activation fail to generalize across contexts

We next asked whether edges uniquely encode belief-inconsistent surprise or whether region of interest (ROI) activation alone predicts surprise across contexts. Following the same leave-one-subject-out model building procedure used to identify the surprise EFPM, we trained the model to identify ROIs in a 268-node whole-brain atlas (Shen et al., 2013) whose activity tracks belief-inconsistent surprise. Unsurprisingly, we found regions whose activation time course tracked belief-inconsistent surprise in the learning task (**Supp Fig. 5**). Regions positively correlated with surprise showed greater activation on trials with more unexpected outcomes (mean within-subject partial *rho*=0.17; two-tailed *p*=1/1001) whereas regions negatively correlated with surprise showed the opposite pattern (mean within-subject partial *rho*=-0.17; two-tailed *p*=1/1001).

However, in contrast to the surprise EFPM, the region-based model failed to generalize to the external dataset (*ß*=-0.022*, t*(65002.919)=-2.415, *p*=0.493). These results suggest that ROI activation alone does not capture belief-inconsistent surprise across contexts. Examining interactions between regions reveals more information about belief-inconsistent surprise.

### The surprise EFPM outperforms predefined brain networks for predicting surprise

External validation demonstrated our model’s sensitivity to belief-inconsistent surprise. Is the surprise EFPM uniquely generalizable or do other brain networks also predict surprise? To address this question, we measured the dynamics of canonical resting-state networks (Finn et al., 2015; **Supp Fig. 3** and **Supp Table 1**) and a validated connectome-based predictive model of sustained attention (Rosenberg et al., 2016a) by averaging the edge time series within each network. We included a network predicting sustained attention because previous work suggests that surprise is attention-orienting (Itti et al., 2009; Yu et al., 2005) and predicts more updates during learning (Nassar et al., 2012).

Several predefined networks fluctuated with belief-inconsistent surprise in the controlled learning task dataset, whereas no canonical networks significantly predicted belief-inconsistent surprise in the NCAA videos. Importantly, these results show that no association between canonical network co-fluctuation and surprise replicate across datasets (**Fig. 4** and **Supp Table 5**).

**Figure 4.**
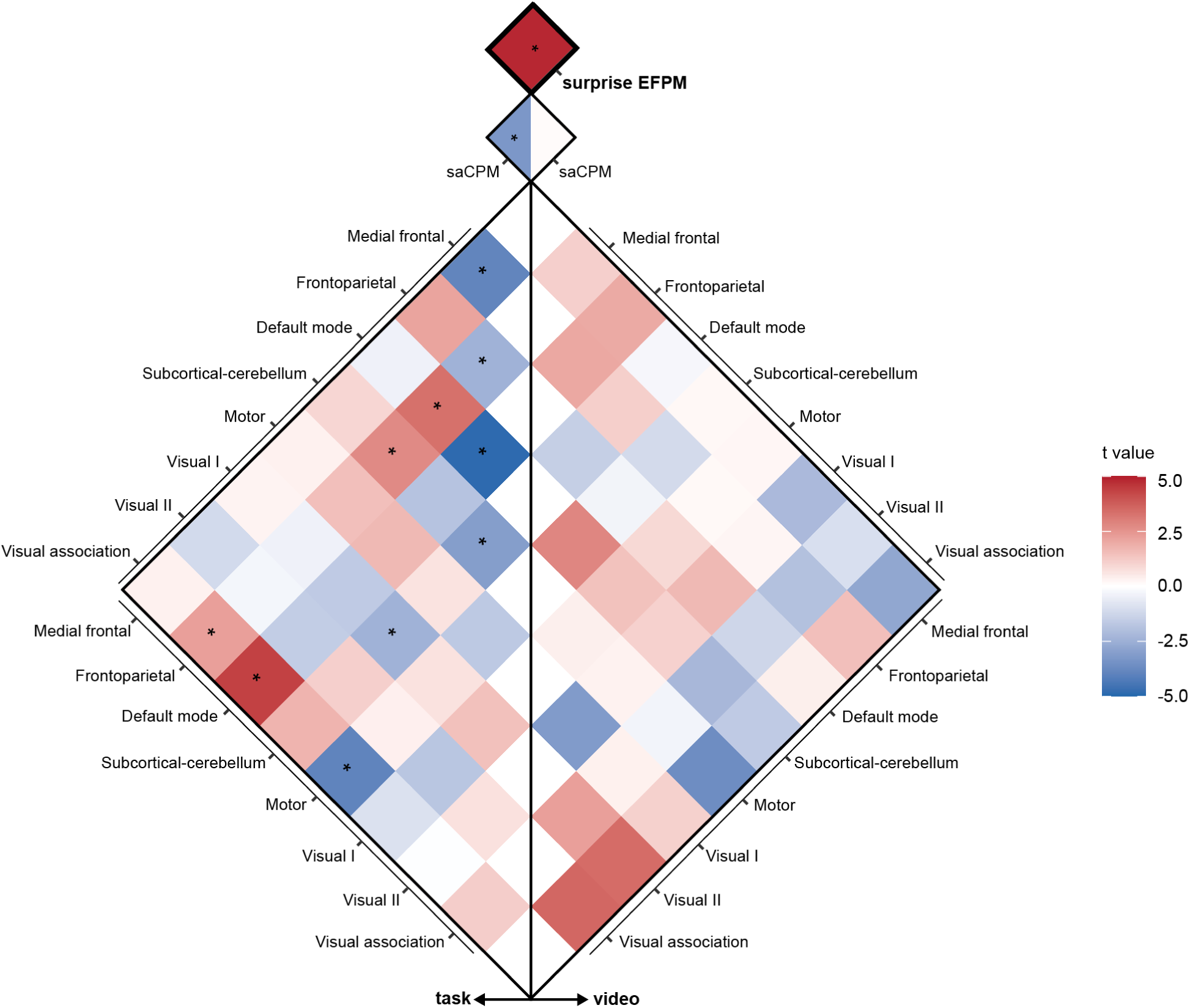
The surprise EFPM defined in the adaptive learning task generalizes to predict belief-inconsistent surprise in a naturalistic viewing context (top square). In contrast, networks and network pairs whose edge time series predicts surprise are not the same across the two datasets (i.e., none of the significant networks on the left and right side of the diamond plot overlap). Significance of the relationship between surprise and each network shown in the plot was assessed by comparing the observed *t* value for the term of the network being tested against a null distribution obtained from running the same linear mixed-effects model with circularly shifted belief-inconsistent surprise time course at a threshold of two-tailed *p*<0.05. Asterisks represent a significant relationship. Positive *t* statistics indicate that a network’s or network pair’s time series is positively correlated with surprise.

While we captured commonalities between surprise across contexts, another interesting question to address is how belief-inconsistent surprise is related to other cognitive processes, like attention and learning. For example, previous work has demonstrated that belief-inconsistent surprise motivates heightened attention to and more incorporation of incoming information (Baldi & Itti, 2010; Itti & Baldi, 2009). We examined this question using a brain network model characterizing sustained attention (saCPM; Rosenberg et al., 2016) to predict belief-inconsistent surprise. The goal of this analysis was to investigate the potential relationship between surprise and sustained attention during task performance and video watching rather than to compare the efficacy of surprise EFPM and saCPM in predicting surprise. Contrary to predictions, the saCPM significantly negatively predicted belief-inconsistent surprise dynamics in the controlled task dataset. In other words, belief-inconsistent surprise was lower when the saCPM predicted that sustained attention was more engaged. Given the important role of surprise in attention and learning, future research could examine relationships between surprise and other components of attention (e.g., selective attention) under a variety of contexts.

### Belief-inconsistent surprise is not predicted by EFPMs built to capture outcome prediction or reward

We next asked whether models built from other factors closely related to surprise—participants’ actual behavioral prediction of the bag-drop location and their affective response to reward—predicted belief-inconsistent surprise in the basketball game.

We trained new EFPMs to predict prediction- and reward-related measures in the adaptive learning task and tested if they generalized to predict belief-inconsistent surprise in the NCAA dataset. EFPMs trained on neither participants’ behavioral prediction of the bag location (*ß*=0.007*, t*(65091.907)=0.700, *p*=0.750) nor the reward (quantified by the number of coins participants caught in their prediction basket on each trial: *ß*=-0.018*, t*(65064.487)=-1.908, *p*=0.145; quantified by the score gained from the coins caught: *ß*=-0.002*, t*(65097.668)=-0.231, *p*=0.874) generalized to predict belief-inconsistent surprise in the NCAA dataset (all *p* values calculated from permutation test via circular shifting). These results suggest that belief-inconsistent surprise in the video was better predicted by an EFPM built on its conceptual counterpart in the learning task, compared to other related task measures.

### The surprise network model is sensitive to violations of psychological expectations

As a final test of model specificity and generalizability, we asked whether the surprise EFPM predicts a different kind of surprise—violations of psychological and physical expectations—in a third independent dataset. We analyzed an openly available fMRI dataset collected while participants watched short video clips showing expected events (expected videos) or violations of expectations (unexpected videos) in scenarios that fall within the domains of psychology and physics (Liu et al., 2024; dataset available at doi:10.18112/openneuro.ds004934.v1.0.0). For example, videos of violations of physics expectations showed solid objects passing through each other or blipping in and out of existence. In videos of psychology expectation violations, there is at least one cartoon agent interacting with an object, another agent, or the environment. Examples includes surprising actions, such as agents changing their goals (e.g., heading in an unpredicted direction) or acting inefficiently (e.g., making an exaggerated move when circumventing an obstacle), and surprising environments, such as agents moving through a solid wall (Saxe et al., 2006).

On each trial, participants viewed a familiarization video clip introducing the scenario (7.75s), followed by either an unexpected or expected video clip (7.75s), a fixation cross (4-10s) and an attention check (2s). To capture brain activity during these clips, we first followed a similar procedure to calculate fMRI BOLD activity and edge time series as those used in the adaptive learning dataset (see Methods). We then calculated the strength of the overlapping network obtained from training on the helicopter task and basketball video datasets (**Fig. 3**). We conducted a paired t-test to compare the two conditions. Demonstrating model generalizability, surprise network strength was higher during unexpected vs. expected psychology-action videos involving agents acting in surprising ways (12 scenarios, with an expected and an unexpected video for each scenario; mean score=.069 vs. –.099; *t*(28)=2.203, *p*=0.036). This relationship, however, was not significant on physics trials involving only physical objects (12 scenarios; mean score=-.019 vs. .050; *t*(28)=-0.852, *p*=0.401) or psychology-environment trials involving interaction between objects and agents (4 scenarios; mean score=.287 vs. .224; *t*(28)=0.578, *p*=0.568). Similarly, Liu et al. (2024) reported neural evidence of domain-specific prediction error in violation of physical but not psychological expectations. Results were similar when controlling for head motion in mixed-effects models.

To test model specificity, we compared the surprise network strength during conditions between which we would not expect surprise to differ: expected psychology vs. expected physics videos. Demonstrating specificity, surprise network strength did not differ between expected psychology and expected physics videos (mean score=.050 vs. 0.03; *t*(28)=0.236, *p*=0.815).

## Discussion

We identified a brain network model, the surprise EFPM, that predicts surprise in controlled and naturalistic tasks from high-frequency edge dynamics. This model generalizes across contexts and outperforms alternative models based on co-fluctuation strength in canonical networks, sliding-window functional connectivity, BOLD activation, and other behavioral measures. Thus, the surprise EFPM captures common neural underpinnings of belief-inconsistent surprise experienced in distinct cognitive contexts in different groups of individuals.

When cognitive constructs such as surprise are studied with different experimental paradigms, an implicit assumption is that a common process or set of processes affects performance across tasks. Evidence for such context-general latent components, however, has been mixed. For example, whereas individual differences analyses of performance on different attention tasks have revealed general attention factors (Huang et al., 2012; Yoo et al., 2022), similar analyses of performance on multiple inhibitory control tasks revealed only low correlations between them (Gärtner & Strobel, 2021). Like attention and inhibitory control, surprise has been assessed with a variety of paradigms hypothesized to evoke similar cognitive and neural responses. These include tasks requiring rare-target detection (Wessel et al., 2016; Mazancieux et al., 2023), conflict detection (Sebastian et al., 2021), and learning in a changing environment (O’Reilly et al., 2013; Meyniel & Dehaene, 2017; McGuire et al., 2014). Surprise has also been assessed in more naturalistic settings, such as during narrative comprehension (Chang et al., 2021; Russo et al., 2020), music listening (Cheung et al., 2019), and sports games viewing (Antony et al., 2021). By demonstrating that edge dynamics associated with belief-inconsistent surprise in one paradigm predict a different measure of belief-inconsistent surprise in two others, EFPM provides evidence for a shared component of surprise across cognitive paradigms. Our results show that we can “translate” belief-inconsistent surprise in different contexts (e.g., controlled tasks vs. naturalistic viewing), operationalized in different ways (e.g., probability of a change point occurring and its uncertainty vs. moment-to-moment change in win probability in a basketball game vs. unexpected vs. expected actions of a cartoon agent) with different classes of models (e.g., reduced Bayesian model quantifying belief vs. categorical summary models) into the common space of the brain.

Using the brain as a common space to understand psychological processes by examining whether multiple paradigms activate the same set of brain regions has been the implicit logic of a subset of fMRI studies since the method was introduced (D’Esposito et al., 1999; Mather et al., 2013). While this approach has generated foundational findings about cognitive theories and brain function, there is no theoretical reason that it need be limited to investigations of regional activation. Here we demonstrated that co-fluctuation in whole-brain data-driven networks—but not regional activation or co-fluctuation in predefined canonical networks—generalized across individuals, tasks and datasets to predict surprise, a result that we may have missed had we built models based on a hypothesis-driven set of features alone. This is in line with the growing evidence that cognitive processes are supported by widely distributed brain networks (Thiebaut de Schotten & Forkel, 2022; Varela et al., 2017). Although these data-driven networks are complex and anatomically extended, including edges within and between cortical, subcortical, and cerebellar regions, they do align with prior work on the neural underpinnings of surprise. Thus, these results underscore the benefits of a data-driven approach for conceptually replicating previously observed brain-behavior associations as well as for generating new hypotheses about networks relevant for behavior.

“Surprise” is broadly used to describe the set of psychological processes involved in expectation violation. These processes include those that precede the experience of expectation violation (e.g., detection of deviants, calculation of prediction error), are maintained throughout the experience (e.g., attention to stimuli, maintenance of an expectation or belief), and follow it (e.g., updating of values, decision policies, and/or beliefs). The surprise EFPM (**Fig. 3**) comprises distributed edges that may support these processes.

First, cross-species work on the neural substrates of prediction error has found evidence that subcortical nuclei for norepinephrine, dopamine, and serotonin are hubs for signaling surprise (Jordan & Keller, 2023; Mazancieux et al., 2023; Valdés-Baizabal et al., 2020). Work has shown that, preceding surprise, deviants in a sequence of stimuli can be signaled by the locus coeruleus (LC) and subthalamic nucleus (STN) regardless of deviation types (Mazancieux et al. 2023; Sebastian et al. 2021). The LC, in particular, is thought to broadcast prediction error signals to cortical regions (Jordan & Keller, 2023). Demonstrating that shared subcortical systems signal surprise induced from different sources, a recent study by Mazancieux et al. (2023) found that LC activity predicts different levels of expectation violation (i.e., locally by oddball stimuli in a short sequence and globally by a rare sequence). Other subcortical areas, including the nucleus accumbens, signal reward prediction error (Rutledge et al. 2010; Schultz et al. 1997; Antony et al. 2021), while the ventral tegmental area signals unsigned prediction error (Barto et al. 2013; Niv et al. 2012; Antony et al. 2021). Taken together, the higher proportion of edges in the subcortical-cerebellar network in the high-surprise compared to the low-surprise network is compatible with observations that subcortical nuclei signal domain-general deviance-detection and prediction errors. However, since the data acquisition was not optimized for subcortical nuclei, this interpretation remains speculative. Future work that maximizes signal-to-noise ratio in subcortical regions in addition to measuring large-scale cortical networks is needed to reveal whether subcortical nuclei activity and broadly distributed edge co-fluctuations explain common or unique variance in surprise in a variety of rich contexts that recruit multiple cognitive processes.

Second, in order to form a general belief about a changing environment, we need to focus attention and maintain expectations (O’Reilly et al. 2013). Using controlled tasks that require participants to explicitly indicate their trial-to-trial predictions, studies have reported that frontoparietal activity (posterior parietal cortex in O’Reilly et al., 2013; McGuire et al., 2014) and functional connectivity (Kao et al. 2020) signal surprise. In line with these findings, the high surprise network included edges in the occipital–parietal and prefrontal-limbic networks (**Fig. 3a**). During naturalistic perception of audiovisual stimuli with rich changes in events, DMN activation has been shown to have higher activation when events are expectation- or schema-inconsistent (Brandman et al., 2021; Dohmatob et al., 2020; Yeshurun et al., 2021) and state transitions in ventromedial prefrontal cortex (vmPFC), a DMN hub, map onto experiences of surprise (Chang et al. 2021). Edges that involve the DMN are present in both of our high and low surprise networks, supporting the hypothesis that the DMN might serve as a hub in high-level prediction-error and belief representations. In contrast to frontoparietal and DMN edges, edges with visual and motor networks were over-represented in the low surprise network, suggesting that greater primary sensory co-fluctuation predicts less higher-level belief updating.

Third, when reality violates our expectation, this discrepancy signals the need to update an existing belief to match with reality. Surprise and updating thus tend to occur close in time. Previous work has found that the anterior cingulate cortex (ACC) and vmPFC are specifically activated during value updating (O’Reilly et al. 2013; Wang et al. 2017; Wu et al., 2021). We observed connections between the limbic system and cortical regions (e.g., prefrontal, parietal and occipital regions) and speculate that the limbic-cortical edges in our high surprise network may reflect higher-level updating processes.

Unlike co-fluctuation dynamics, neither regional activity dynamics nor sliding-window functional connectivity dynamics generalized across datasets to predict surprise. The lack of generalizability of regional activity dynamics demonstrates the unique predictive power of region interactions rather than region activity alone. We speculate that this is because activity in individual regions may capture task-specific information unique to the training set, which does not predict the belief-inconsistent surprise signal when the context changes. This hypothesis aligns with recent observations that higher-order interaction dynamics capture higher-order cognition dynamics, whereas activation patterns predict lower-level sensory information (Owen et al., 2021). The lack of both within- and across-dataset prediction from sliding-window functional connectivity dynamics suggests that models built from higher-frequency edge co-fluctuations better capture fine-grained changes in cognitive processes. Co-fluctuation circumvents the loss of information that can result from smoothing transient activities within a window to calculate regional interactions and may thus be a better choice for predicting moment-to-moment fluctuations in other cognitive processes beyond surprise.

Both sensitivity (true positive rate) and specificity (true negative rate) provide crucial information about a predictive model’s performance. Brain-based predictive modeling research tends to focus on the former, evaluating models’ sensitivity by testing them with held-out and/or out-of-sample data (Cao et al., 2019; Rosenberg & Finn, 2022; Woo et al., 2017). Here we assessed both aspects: we demonstrated the model’s sensitivity to surprise by training it on controlled task data and testing it on naturalistic data acquired when a different group of individuals performed a distinct task operationalizing a similar psychological process and vice versa. We assessed model specificity by demonstrating that the EFPM built from helicopter task data predicted surprise during basketball games even when controlling for other factors (e.g., amount of time remaining, team in possession of the ball, auditory envelope, luminance). In contrast, models trained to predict closely related task measures (behavioral prediction and reward) did not predict belief-inconsistent surprise in the basketball games. Furthermore, the EFPM trained on video data predicted belief-inconsistent surprise, but not behavioral prediction or reward received, in the helicopter task. As a final test of sensitivity in a third dataset, we showed that the surprise model distinguished videos of cartoon agents that did vs. did not violate expectations. Demonstrating specificity, the model did not distinguish different types of videos that did not violate expectations. Thus, the surprise EFMP is a relatively sensitive and specific predictor of surprise.

We point out a few limitations of our work. First, the amount of variance that can be explained by the model is relatively small. A robust and reliable predictive model, however, need not achieve a particular accuracy threshold to be theoretically meaningful. Our cross-dataset sensitivity and specificity tests suggest that some aspects of surprise are domain-general rather than domain-specific—a result that we could not have observed in these data with behavioral measures or computational models alone. Second, the surprise network generalized to predict expectation violations involving the actions of cartoon agents, but not agents interacting with the environment or physical objects alone. A speculative explanation for this lack of generalization comes from studies showing that both prediction and prediction errors about physics events may recruit domain-specific regions in the brain (Fischer et al. 2016; Liu et al. 2024), although this remains to be tested in further work. Finally, the surprise EFPM is a highly extended data-driven network which can impede anatomical interpretability. Identifying edges that consistently predicted surprise in two datasets helped constraint networks to those most reliably predictive, and computational lesioning analyses helped identify edges on which the model most strongly relied. Furthermore, models based on co-fluctuations within and between canonical networks alone did not reliably predict surprise, underscoring the importance of whole-brain prediction approaches for building generalizable models of surprise. That said, it will be important for future work to explore different model-building algorithms and feature-selection approaches and thresholds to identify the necessary and sufficient set of edges in a maximally generalizable model of surprise.

Looking ahead, the surprise EFPM motivates further research on both surprise and dynamic predictive modeling. First, the surprise EFPM itself can be validated using more measures, including subjective measures (e.g., facial expressions or densely sampled subjective ratings of surprise), outputs of models that predict behaviorally relevant outcomes from audiovisual, semantic, or linguistic input (e.g., large-language-model-generated unsigned prediction errors), or other neural signatures of expectation violation (e.g., LC activation). Furthermore, work could use this model to study the interaction between surprise and other cognitive processes such as aspects of attention. When we applied an existing brain network model of sustained attention, the saCPM, to edge co-fluctuation data, we found a negative relationship between surprise in the task and participants’ signature of sustained attention. Although we originally hypothesized that surprise would be associated with increased sustained attention, these results suggest that surprise may break participants’ sustained period of focus by introducing changes to the environment. Future research could answer open questions about the relationship between surprise and attention dynamics, such as, whether surprise captures, orients, and enhances *selective* attention while interrupting states of *sustained* attentional focus.

Second, EFPM is a general framework that can be applied to predict other cognitive, attentional, and affective processes and states from neuroimaging data. This approach offers new perspectives to analyze data from existing cognitive paradigms and possibilities for identifying signatures of cognitive states fluctuations from large-scale brain networks (Flavell et al., 2022; Greene et al., 2023; McCormick et al., 2020). Developments in open-data practices including code, data, and model sharing facilitate model sensitivity and specificity testing with data acquired at multiple sites with a range of tasks and participant populations. After establishing a shared component through predictive mapping, research could then generate hypotheses about what accounts for the generalizability and predictive boundaries of the model, and design follow-up studies to test these hypotheses. For example, future research could use this framework to capture other task-induced states (e.g., affective state, attentional state, and working memory processes) and then use out-of-sample generalization compare and contrast across paradigms, providing insights into the extent a set of paradigms recruit common underlying processes.

Now reconsider the question raised at the beginning: are there shared processes in how our brain responds to unexpected changes in surprising moments—be it a surprise party, a lab task, or a suspenseful basketball game—despite being in completely different contexts? In crossing traditional cognitive paradigms, predictive modeling based on edge dynamics provides new evidence that there are.

## Data and Code Availability

Helicopter learning task data are available at https://openneuro.org/datasets/ds003772/versions/1.0.1 (McGuire et al., 2021). NCAA viewing data are available at https://openneuro.org/datasets/ds003338/versions/1.1.0 and https://dataspace.princeton.edu/handle/88435/dsp019k41zh56d (Antony et al., 2021). Violation of expectation (VoE) data are available at https://openneuro.org/datasets/ds004934/versions/1.0.0 (Liu et al., 2024). Analysis code associated with this paper, including code for training and testing new edge-fluctuation-based predictive models, is available at https://github.com/ZiweiZhang0304/Surprise_EFPM.git. We used the following versions of the softwares: MATLAB (R2020a), AFNI (v.19.0), R (version 4.0.5), and Python (3.12.2).

## Acknowledgements.

We thank Joseph T. McGuire, Joseph W. Kable, and Chang-Hao Kao for sharing and addressing questions about the MRI and behavioral data associated with McGuire et al. (2014) and Kao et al. (2020). We thank James W. Antony for sharing and addressing questions about the MRI data, behavioral data, and analysis scripts associated with Antony et al. (2021). We thank Shari Liu for sharing and addressing questions about the MRI data associated with Liu et al. (2024). We also thank Joshua Faskowitz and colleagues for making the code associated with Faskowitz et al. (2020) publicly available, which we adapted and used in this work. We thank Veritone and the NCAA for permission to use basketball game screenshots. We thank Yuan Chang Leong, Kwangsun Yoo, Hayoung Song, and William X. Q. Ngiam for helpful feedback and suggestions on this work. Research was supported by National Science Foundation BCS-2043740 to M.D.R. and resources provided by the University of Chicago Research Computing Center.

## Online Methods

### Adaptive learning task dataset

The raw MRI data associated with the adaptive learning task (McGuire et al., 2014) was downloaded from OpenNeuro (https://openneuro.org/datasets/ds003772/versions/1.0.1), and the output from the reduced Bayesian model was shared by Kao et al. (2020) and McGuire et al. (2014).

#### Participants

The dataset was described in detail in previous work (McGuire et al., 2014). Participants were recruited from the University of Pennsylvania community: *n* = 32, 17 female, mean age = 22.4 years (SD = 3.0; range 18–30). Human subject protocols were approved by the University of Pennsylvania Internal Review Board. Informed consent was provided by all participants.

#### Learning task

In the scanner, participants performed an adaptive learning task in which they had to predict the location of an upcoming object, a dropped bag (**Fig. 1a**). The location of the object was drawn from a distribution hidden to the participants, with a certain mean and standard deviation. The hidden distribution remained unchanged most of the time but changed unexpectedly (probability of change = 0.125) where the mean of the distribution was re-drawn from a uniform distribution [0-300]. The outcome was then revealed by the falling coins into the basket. The moment when an outcome that is far from the predicted location (i.e., where the participants placed the basket) is revealed could be considered as a surprising moment. A random manipulation of reward was also implemented, where on each trial the value of the coins was randomly determined to be positive (yellow coins) or neutral (gray coins). The participants’ scores were determined by the number of coins caught with positive values but coins of both colors were equally informative for making future predictions. At the start of each trial, participants needed to maneuver the joystick to the ‘home position’ located on the right-hand side of the screen to retrieve the bucket, after which they could relocate it to make predictions. Participants performed four 80-trial practice runs before MRI scanning, where they saw the helicopter representing the generative distribution, were informed about the change point, and that the best strategy to perform the task is to keep track of the helicopter. These instructions facilitated participants to form beliefs about the helicopter location. McGuire et al. (2014) found a relationship between participants’ behavioral learning rate and the normative learning rate, suggesting that the surprise information modulated participants’ behavior.

#### Normative model of task surprise

McGuire et al. (2014) developed a reduced Bayesian model to operationalize the amount of surprise in the environment. This normative model generates a trial-to-trial estimate of two measures: change-point probability (CPP), which describes the probability of a change point (i.e., a mean shift in the generative distribution) occurring and relative uncertainty (RU), which describes the uncertainty of this change truly occurring relative to noise from the variance of the two distributions. What should govern how much participants update their belief based on what they learn from the prediction error (prediction-outcome) on a trial-to-trial basis? Bayesian surprise measure scales the amount of belief updating from prediction error. Highly surprising outcomes might signal changes in the environment and thus render belief updating and learning important. A learning rate was calculated in Nassar (2010) based on these two measures:

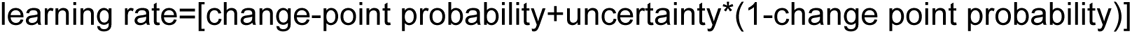

Belief on each trial after outcome was then updated using a delta updated rule, scaled by the learning rate that is constrained between zero and one. In previous work, Nassar et al. (2010) has demonstrated that the learning rate term in this parsimonious reduced Bayesian normative learning model efficiently tracked human updating behavior. We used this learning rate as a measure of belief-inconsistent surprise because CPP and RU scales belief updates in the learning process.

#### MRI data acquisition

Data acquisition was described in McGuire et al. (2014). MRI data were gathered using a 3T Siemens Trio machine with a 32-channel head coil. Gradient-echo imaging (EPI) was used to acquire functional data, with an isotropic voxel size of 3 mm, a 64 x 64 matrix, 42 tilted axial slices, a TE of 25 ms, a flip angle of 75°, and a TR of 2500 ms. The study consisted of four runs, each with 226 images. T1-weighted MPRAGE structural images were also obtained, with a voxel size of 0.9375 x 0.9375 x 1 mm, a 192 x 256 matrix, 160 axial slices, a TI of 1100 ms, a TE of 3.11 ms, a flip angle of 15°, and a TR of 1630 ms. Additionally, matched field map images were collected with TE of 2.69 and 5.27 ms, a flip angle of 60°, and TR of 1000 ms.

### Naturalistic sports viewing dataset

The MRI data associated with NCAA sports viewing (Antony et al., 2021) was obtained from https://openneuro.org/datasets/ds003338/versions/1.1.0. The non-MRI data containing all video information and surprise model output were obtained from https://dataspace.princeton.edu/handle/88435/dsp019k41zh56d.

#### Participants

The dataset was described in detail in previous work (Antony et al., 2021). Twenty participants (6 female, 18-35 years old) fluent in English with normal or corrected-to-normal vision were recruited. Written informed consent was obtained in a manner approved by the Princeton University Institutional Review Board.

#### Task

The last five minutes from all 32 Round-of-64 2012 men’s NCAA® tournament games were acquired as audiovisual files for a fee and with permission from Wazee Digital. In the scanner, participants watched nine of those videos (**Fig. 1**). In each of the three MRI runs (mean run length=19.43 mins, mean game length=6.48 mins), participants alternated between game viewing and free recall phases.

#### Model of naturalistic surprise

To quantify surprise, Antony et al. (2021) first created a 4-factor model to predict the win probability of a given team based on four factors: the difference in score between the teams oriented to be positive when the ‘‘home’’ team is winning and negative when they are losing, the amount of time remaining, which team is in possession of the ball, and, where publicly available, an adjustment based on team strength. Specifically, the game situation on the court was categorized into bins considering every score difference ranging from -20 to +20 for the home and visitor team scores, each time window within 6 seconds (e.g., 60-54 seconds left in the game), and the possession of either the home or visiting team.

From the game results in the entire season, one could first determine the relative strength of a given pair of teams by calculating the expected amount that the home team should outscore the away team over an entire game. However, this expected score difference does not remain the same across the duration of the game since with less time remaining it is less possible to produce a score that deviates from the expected difference. The degree to which the home team could out-score the away team thus also depends on the time remaining in the game at a given time point. Consider an example: one might first find that the home team is expected to outperform the away team by a score of six. Now if it is half-way into the game, the home team will only be expected to out-score the away team by three. If the observed score difference at the current time point is one (home team losing), they will be expected to out-score the away team by only two points by the end of the game. Finally, the win probability might also depend on which team is currently in possession of the ball. When implementing the model to generate predictions about win probability, the authors created a “look-up table” for each combination of the bins of these factors and calculated the percentage of games that the home team ended up winning. A “look-up table” with three factors was created: expected score difference by the end of the game (current score difference – potential difference and thus combined the first two factors of the four-factor model), time remaining, possession of the ball (for more detailed information on this model, see Antony et al., 2023). Moment-to-moment win probability of the nine games were then “read out” from locating all the games that had the factors matching with the three input factors, and then calculating how often the home team won among these games.

All regular season men’s basketball games from the 2012 season were used to train the model. The data was first split into half for cross-validation. The 4-factor model was trained using half of the data and then validated by predicting the win probabilities in the left-out half and comparing the predictions with that from an expert sports analyst (https://www. kenpom.com/).

The measure of surprise reflects how belief in some outcome (e.g., which team will win) changes over time. Surprise was calculated as the absolute value of the change in the win probability time course at each possession boundary.

The change in win probability at a given time point relative to the previous point was further separated into belief-consistent and belief-inconsistent surprise. The authors first identified moments when the home team is winning if the win probability metrics calculated based on the 4-factor model were greater than 50%. Then only a decrease in the win probability metrics would result in non-zero belief-inconsistent surprise. Similarly, moments when the home team was losing were identified if the win probability is less than or equal to 50% and belief-inconsistent surprise is non-zero only when there is an increase in the win probability. For all moments with a non-zero belief-inconsistent surprise, the value is numerically equal to surprise (i.e., the absolute difference in win probability between the current and previous time point).

Antony et al. (2021) demonstrated that belief-inconsistent surprise correlated with participants’ subjective segmentation of event boundaries, and that segmentation was more robust when the situations in the basketball games violate the predominant belief.

### Violations of psychological and physical expectations dataset

We reanalyzed MRI data associated with experiment 2 in the violations of psychological and physical expectations (VoE) dataset (Liu et al., 2024). Data were obtained from https://openneuro.org/datasets/ds004934/versions/1.0.0. Data from experiment 2 were analyzed because it included more stimuli and participants than experiment 1.

#### Participants

The dataset was described in detail in previous work (Liu et al., 2024). In Experiment 2, 33 participants (Mean age = 25.7y, range 18-45; 30 right-handed; 21 female, 12 male; 19 White; 14 Black, Asian, Latine, or multiracial) were recruited. Twenty-nine participants gave consent to share their fMRI data and we thus analyzed data from these 29. Written informed consent was obtained in a manner approved by the MIT Committee on the Use of Humans as Experimental Subjects.

#### Task

Participants underwent 4 runs of a violations-of-expectations task. On each trial, participants viewed a familiarization video clip introducing the scenario (7.75s), followed by either an unexpected or expected video clip (7.75s), a fixation cross (4-10s) and an attention check (2s). For the attention checks, participants were instructed to press a button if the fixation cross was the letter X instead of a plus symbol (+) (33% of trials). There were 12 scenarios from the domain of physics, where inanimate objects were shown (e.g., surprising clips involved solid objects passing through each other). Similarly, there were 4 scenarios from the domain of psychology-environment, where humanoid cartoon agents interacted with objects in the environment (e.g., surprising clips involved an agent passing through a solid wall). There were 12 scenarios from the domain of psychology-action, where these cartoon agents engaged in actions (e.g., surprising clips involved an agent changing goals or acted inefficiently). Trials on which participants failed the attention check by making an error (number and percentage of excluded trials=178 and 9.59%) were excluded from the analysis. Video stimuli are available at https://osf.io/sa7jy/.

#### MRI data acquisition

Data acquisition is described in Liu et al. (2024). Neuroimaging data were acquired using a 3-Tesla Siemens Magnetom Prisma scanner with a 32-channel head coil. Ten runs of functional data were collected (gradient-echo EPI sequence sensitive to Blood Oxygenation Level Dependent (BOLD) contrast in 3mm isotropic voxels in 50 interleaved near-axial slices covering the whole brain; EPI factor=70; TR=2s; TE=30.0ms; flip angle=90 degrees; FOV=210mm).

### MRI data preprocessing

We applied the same preprocessing steps to all MRI datasets using the AFNI software (v.19.0 Cox, 1996, 2012). afni_proc.py was used to run preprocessing steps. Outliers in functional data were first removed with despiking (*despike*), corrected for slice timing (*tshift*), aligned with anatomical scan (*align*), and then registered to the MNI space (*tlrc*). Volume registration (*volreg*) was also performed where the functional volumes were registered to the volume that was considered a minimum outlier. We also regressed out the covariates of no interest from the data by including the following regressors: mean signal from two eroded masks (ventricles and white matter); global signal (Faskowitz et al., 2020; Rasero et al., 2022); a 24-parameter motion model (6 motion parameters, 6 temporal derivatives, and their squares). We defined *a priori* to censor volumes where more than 10% of voxel were considered outliers, and volumes with head motion parameter derivatives’ Euclidean norm surpassing 0.25 from the time-series.

To obtain the mean time series per ROI, Shen 268-node functional brain atlas (Shen et al., 2013) was used to parcellate the voxels into 268 nodes (*3dNetCorr*). This atlas was defined using a groupwise graph-theory-based parcellation algorithm that maximized the similarity of the timeseries of the voxels within each node. Data were resampled to MNI space with a resolution of 3 × 3 × 3 (*3dresample*). We also defined *a priori* that nodes missing in each subject will be removed from all subjects. This process resulted in the removal of five nodes from the adaptive learning task data (two nodes from right temporal and three nodes from left temporal regions) and the naturalistic sports viewing data (three nodes from right cerebellum, one node from left temporal, and one node from left cerebellum regions), respectively. For the naturalistic sports viewing dataset, data from the first 10 s (TRs) of each video were removed to avoid the effects of strong onset responses (Antony et al., 2021; Nastase et al., 2019). We then aligned the data for the nine games in the same order within each participant in preparation for the calculation of the co-fluctuations described in the section below.

To control for head motion, we also calculated the framewise displacement (FD) as follows:

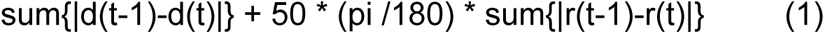

where d denotes translation distances {x,y,z}, and r denotes rotation angles {α,β,γ} (Patel et al., 2014). In the adaptive learning dataset, displacement was calculated using the same average approach as trial-to-trial brain activity (see the section titled *Adaptive learning task dataset*. in Methods.). Specifically, we located the TR of outcome onset (the i^th^ TR) and then took the average of the (i+2)^th^, (i+3)^th^, (i+4)^th^ TRs for each of the six motion parameters {x,y,z,α,β,γ}. After getting the peak of the motion effects, we calculated the framewise displacement quantifying how much the head has moved following Equation 1. In the naturalistic sports viewing dataset, FD was calculated for every TR with the above equation.

#### Calculating edge time series with co-fluctuation

We quantified the extent to which a pair of nodes in the brain are deflected in the same direction at every TR following the edge-time-series method developed by Faskowitz et al. (2020; code adapted from: https://github.com/brain-networks/edge-centric_demo/blob/main/fcn/fcn_edgets.m). The time series of each edge is defined as the element-wise product of the *z*-scored time series in a pair of nodes in the brain. Each time point in the edge time series quantifies the extent to which the activity in the two regions at that moment in time deflects in the same direction. For example, if two nodes both show above baseline activation at precisely the same moment in time, the value in the edge time series is positive. Similarly, a time point in the edge time series is negative when one region is activated and the other one is deactivated, and is close to zero when activities in both regions are close to baseline. The calculation of edge time series can thus be viewed as an intermediate step to calculating Pearson correlation, a standard way to measure functional connectivity or the statistical interdependency between two regions. While calculating Pearson correlation involves three steps: *z*-scoring time series between two regions, taking the elementwise product of the two, and calculating the mean of this product to obtain a single value across time, the calculation for edge-time-series simply involves the first two steps, thus resulting in a value at each time point.

#### Adaptive learning task dataset

We extracted trial-to-trial brain activity in each of the 268 brain nodes from the preprocessed time series data by first locating the TR of the outcome onset of the learning trial (the *i*^th^ TR) and then taking the average of TRs *i*+2, *i*+3, *i*+4. This calculation approximately captures the peak of the corresponding brain activities in the TR of interest. The time series for each run and each participant were first *z*-scored within each region and the time-resolved co-fluctuations were then calculated as the vectors of element-wise products for every pair of regions (Faskowitz et al., 2020), which resulted in a 120 (trials) by 35,778 (pairs of regions) edge time series matrix for each run and each participant.

#### Naturalistic sports viewing dataset

Since naturalistic viewing is continuous, we computed co-fluctuations at every TR. We *z*-scored the time series for each participant within each region and then calculated co-fluctuations following the same method above, namely, as the vectors of element-wise products for every pair of regions. This calculation resulted in a 3,408 (TRs) by 35,778 (pairs of regions) edge time series matrix for each participant.

#### Violations of psychological and physical expectations dataset

We first followed the same procedures to calculate fMRI BOLD activities and edge time series as those used in the adaptive learning dataset (see *Adaptive learning task dataset* above). Specifically, to capture edge strength in each video clip, we marked the start of each video (*i^th^* TR) and averaged co-fluctuation values across TRs (i+2)^th^, (i+3)^th^, (i+4)^th^, (i+5)^th^, (i+6)^th^ to capture the duration of the video.

#### Pre-defined networks and their co-fluctuation strength

To assess the spatial specificity of the surprise EFPM, we asked whether the strength of networks that were previously defined to predict a variety of cognitive task performance vary with belief-inconsistent surprise.

The first set of networks we used is the sustained attention connectome-based predictive model (saCPM), which contains a set of functional brain networks whose strength during both a sustained attention task and rest predicts individual differences in performance (Rosenberg et al., 2016a) as well as how sustained attentional states vary within individuals over time (Rosenberg et al., 2020). Strength of saCPM also generalized to predict sustained attention task behavior in previously unseen individuals. Specifically, the saCPM contained two networks: a high-attention network, whose strength positively correlated with sustained attention task performance, and a low-attention network that showed the opposite relationship. The high-attention network comprised 757 edges and the low-attention network, 630 edges. The model is available at https://github.com/monicadrosenberg/Rosenberg_PNAS2020.

To obtain a moment-to-moment measure of saCPM strength, we multiplied the mask for positive and negative networks (where the masks contain values of ones and zeros to indicate which edges are in each of the two networks) separately with the edge time series matrix. We then calculated the mean co-fluctuation value of the edges within the positive and negative network, respectively. The total saCPM network strength on each trial was then calculated as the difference between the two: positive network strength – negative network strength.

The next set of networks we used was defined by Finn et al. (2015) from the same set of 268 nodes used here (Shen et al., 2013). The 268 nodes were defined into eight clusters representing the following networks: medial frontal (MF), frontoparietal (FP), default mode (DM), subcortical/cerebellum (SubcortCere), motor (MT), visual I (VI), visual II (VII), visual association (VA). For each one of the eight networks, we applied the mask denoting which node belongs to the current network to the edge-time-series matrix and calculated the mean co-fluctuation of the edges within this network to obtain TR-by-TR time series of the canonical network strength.

### Building an edge-fluctuation-based predictive model (EFPM) of belief-inconsistent surprise in the adaptive learning task dataset

#### Internal cross-validation of surprise EFPM

We used leave-one-out cross-validation (LOOCV) within the controlled learning dataset to select edges that meaningfully correlate with belief-inconsistent surprise. Specifically, in each fold of validation, we selected 31 (i.e., *n*-1) participants as the training set and then calculated the partial Spearman correlation (*rho*) between belief-inconsistent surprise and the co-fluctuation time series of every edge in the brain, controlling for head motion (i.e., framewise displacement as described in the above *MRI data preprocessing section*). We performed a Fisher *z*-transformation on the partial correlations (*rho*), followed by a one-sample t-test of these 31 *rho* values against a null value of zero. We then identified edges that exhibited statistically significant positive and negative correlations with belief-inconsistent surprise across the group, based on a significance threshold of *p*<0.05. Edges positively correlated with belief-inconsistent surprise after thresholding was selected into the positive network, while the edges that are negatively correlated with belief-inconsistent surprise with the same thresholding criteria were selected into the negative network.

After each training fold is finished, the positive and negative networks were applied to the edge time series of the left-out participant to obtain the trial-to-trial strength of these selected edges. Specifically, we separately averaged the edge time series in the positive and negative surprise networks to obtain a single network time series for each. We again calculated the *rho* between the held-out participant’s belief-inconsistent surprise and the strength of the network identified from the training participants, controlling for head motion. We repeated this process 32 (i.e., number of participants in the controlled learning task) times, to obtain the observed *rho* values.

#### Significance testing and surprise EFPM network definition

To assess the significance of the edges identified from this process, we iterated the LOOCV edge selection process 1000 times using phase randomized network score within each subject and averaged across subjects to obtain a null distribution. Phase-randomization was implemented using function: https://github.com/OHBA-analysis/MEG-ROI-nets/blob/master/%2BROInets/generate_phase_su rrogates.m; Colclough et al., 2015. We then calculated the non-parametric *p* value as follows:

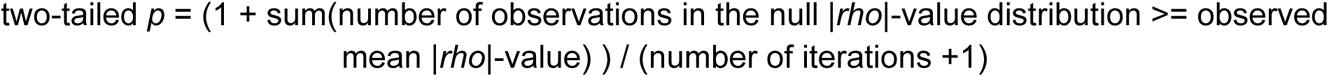

and used a threshold of *p*<0.05 to determine the significance of the edges selected.

The final surprise EFPM contained two networks: a high-surprise network and a low-surprise network. The high-surprise network contained edges that are positively correlated with belief-inconsistent surprise in every one of the 32 cross-validation folds, while the low-surprise network contained edges that are negatively correlated with belief-inconsistent surprise in all folds.

In addition to the approach we used to obtain the null distribution described above, we also obtained another null distribution by randomly selecting same-size edge sets in each training fold for prediction and repeated this 1000 times.

#### Testing external generalizability of the surprise EFPM to the naturalistic sports viewing dataset

Next we built a linear mixed-effects model to see whether surprise EFPM strength, along with other regressors that we are controlling for in the video, predicts surprise in a naturalistic context. To obtain surprise EFPM scores, we multiplied the mask for the high and low surprise EFPM (where the masks contain values of ones and zeros to indicate which edges are in each of the two networks) separately with the edge time series matrix. We then calculated the mean co-fluctuation value of the edges within the high and low surprise networks, respectively. The total surprise EFPM score for each participant at each time point was then calculated as the difference between the two: high surprise network score – low surprise network score.

Measures of surprise, along with other variables in the video, were convolved with a canonical hemodynamic response function and *z*-scored within participant before they were entered as the predictors in the following linear mixed-effects model:

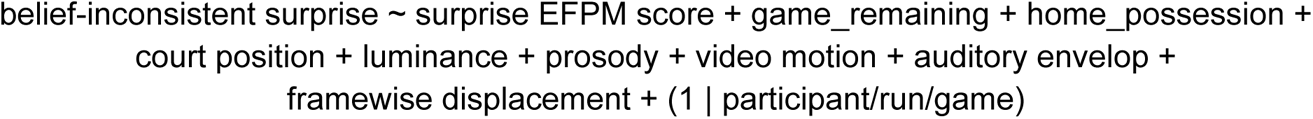

where we included a set of additional regressors (Antony et al., 2021), including the amount of game remaining in seconds of game time at the start of the possession; which team was in possession of the ball; the auditory envelope; global video luminance; global video motion; commentator prosody; court position of the ball (left or right); framewise displacement of the head. The model was run using the lmerTest (3.1-3) package in R (version 4.0.5).

#### Significance testing

We ran the above model repeatedly by circularly shifting both the forward and backward belief-inconsistent surprise time series within each game, with the restriction that any given TR cannot fall within the nearby five TRs after circular shifting. A null distribution of the model *t*-value was obtained from these iterations and the observed *t* value of the surprise EFPM term was compared against the null distribution. We then calculated the *p* value as follows:

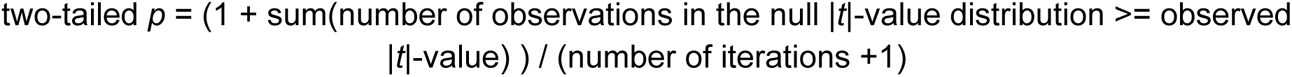

and used a threshold of *p*<0.05 to determine the significance of the generalizability of the surprise EFPM.

#### Predicting belief-inconsistent surprise with sliding-window functional connectivity and testing its generalizability

Dynamic functional connectivity was calculated with 3 different window sizes (11, 21, 31 TRs, corresponding to 27.5, 52.5, and 77.5 seconds) in the helicopter task and basketball video datasets. Following previous work (Allen et al., 2014; Song et al., 2021), we calculated Fisher’s z-transformed Pearson’s correlation between the time courses of every region-pair with a tapered sliding window (code adapted from Song et al. (2021); https://github.com/hyssong/NarrativeEngagement). The peak correlation at surprise onset (i^th^ TR) was then captured by the (i+3)^th^ TR to account for hemodynamic delay. These resulted in *n* (trials) by 35,778 (pairs of regions) edge time series matrix for each run and each participant, where *n*=120,114,108 for window=11,21,31 TRs. These matrices were then used as input to the same EFPM training pipeline used to identify an edge-co-fluctuation-based model.

We also tested the generalizability of the model built from the sliding window approach in the adaptive learning task to the NCAA dataset. To do so, we identified the predictive networks in the training set and calculated their score in the test data following the same method as above (see *Testing external generalizability of the surprise EFPM to the naturalistic sports viewing dataset* in *Methods*). Significance was also examined with the same circular shifting procedure.

#### Testing external generalizability of the overlapping surprise EFPM to the VoE dataset

We calculated the strength of the overlap surprise network (**Fig 3**.) during expected vs. unexpected events in the VoE dataset following the same procedure as the learning task (see *Adaptive learning task datase*t in *Methods*). After obtaining a strength of the surprise network for each of the unexpected and expected clips, we z-scored the surprise EFPM strength values within each participant and calculated a mean strength for each one of the two events (expected and unexpected) in each participant. We then conducted a paired t-test comparing the strengths under the two conditions.

#### Computational lesioning

We performed a computational lesioning analysis to further understand the anatomy of the surprise EFPM. If a predefined network plays an important role in predicting belief-inconsistent surprise, we would expect that lesioning that network would lead to failure in generalization to the basketball-viewing dataset. To perform this analysis, for each network, we first excluded all edges that involve at least one node in this network from both the high and low surprise networks in the surprise EFPM, and then calculated the score of the remaining edges in the same way we calculated the network strength for the intact surprise EFPM. The network strength for each one of the lesioned EFPM were then separately entered as input to the linear mixed model used to assess the surprise EFPM generalization (see *Testing external generalizability of the surprise EFPM to the naturalistic sports viewing dataset*). Assessment of statistical significance was consistent with that of the surprise EFPM.

#### Comparing the generalizability of the surprise EFPM with that of the predefined networks

Next, we asked whether the network strengths calculated from co-fluctuations within predefined networks also varied with belief-inconsistent surprise within each dataset and consistently across datasets.

Specifically, the predefined networks we used are the eight functional networks used in Finn et al. (2015) and the sustained attention connectome-based predictive model (saCPM) developed by Rosenberg et al. (2016a). We then calculated the strength of co-fluctuations in each of the predefined networks by applying a mask of each network to the edge time series matrix and then taking the average of the values remaining in the masked edge time series.

First, to ask whether the strength of co-fluctuations within each of the pre-defined networks can predict the belief-inconsistent surprise in the adaptive learning task, we built a mixed-effects model specified as follows:

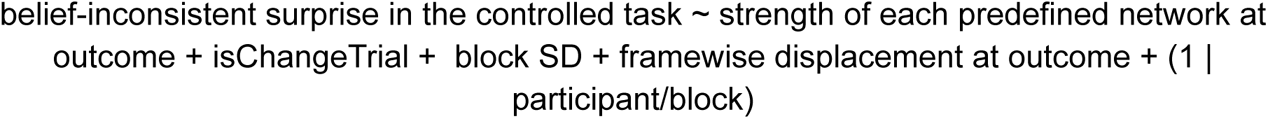

For each of the eight networks, the regressors in the model included the strength of the predefined network at outcome, whether a trial is a change trial (factor: 0 vs. 1), the standard deviation within a block (factor: 10 vs. 25) and framewise displacement as fixed effects. The numerical regressors, strength of the networks and framewise displacement, were *z*-scored at outcome before entering them into the model to transform them to the same scale for model interpretation. We also included participant and block (factor: 1-4) as random effects with varying intercepts.

For the same process in the naturalistic sports viewing dataset, we ran a model specified below (similar to the model used in the *Testing external generalizability of the surprise EFPM to the naturalistic sports viewing dataset* section):

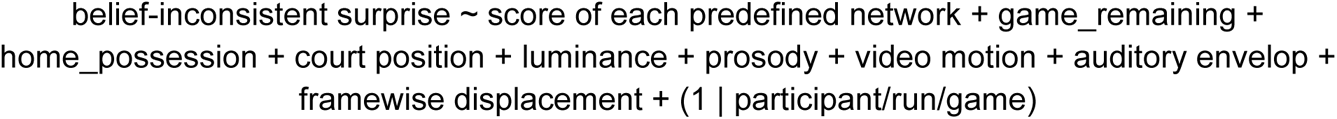

where we only changed one of the regressors from the strength of the surprise EFPM to the strength of each predefined network. Significance was assessed as described in the *Testing external generalizability of the surprise EFPM to the naturalistic sports viewing dataset* section.

#### Predicting belief-inconsistent surprise with region-level BOLD activation

Using the same EFPM approach, but this time applied to BOLD signal rather than edge time series, we trained a model to predict surprise from ROI time series. This was accomplished using a matrix of dimensions 268 rows by the number of TRs, in contrast to the previous matrix size of 35778 rows by the number of TRs. Generalizability to the naturalistic sports viewing dataset was assessed with the model below:

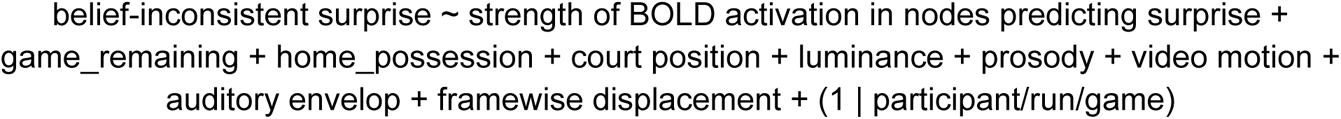

where the only term that was different from the EFPM is that instead of using the strength of the surprise EFPM, here we used the strength of BOLD activation in nodes we identified as correlating with belief-inconsistent surprise in the adaptive learning task dataset based on their activities. Significance was assessed as described in the *Testing external generalizability of the surprise EFPM to the naturalistic sports viewing dataset* section.

## Notes

https://openneuro.org/datasets/ds003772/versions/1.0.1

https://openneuro.org/datasets/ds003338/versions/1.1.0

https://openneuro.org/datasets/ds004934/versions/1.0.0

### Competing Interest Statement

The authors have declared no competing interest.

### Summary of Updates

Figure 1 revised; Figure 2 revised

