## Supplemental material for "Brain network dynamics predict moments of surprise across contexts"

### Supplementary information

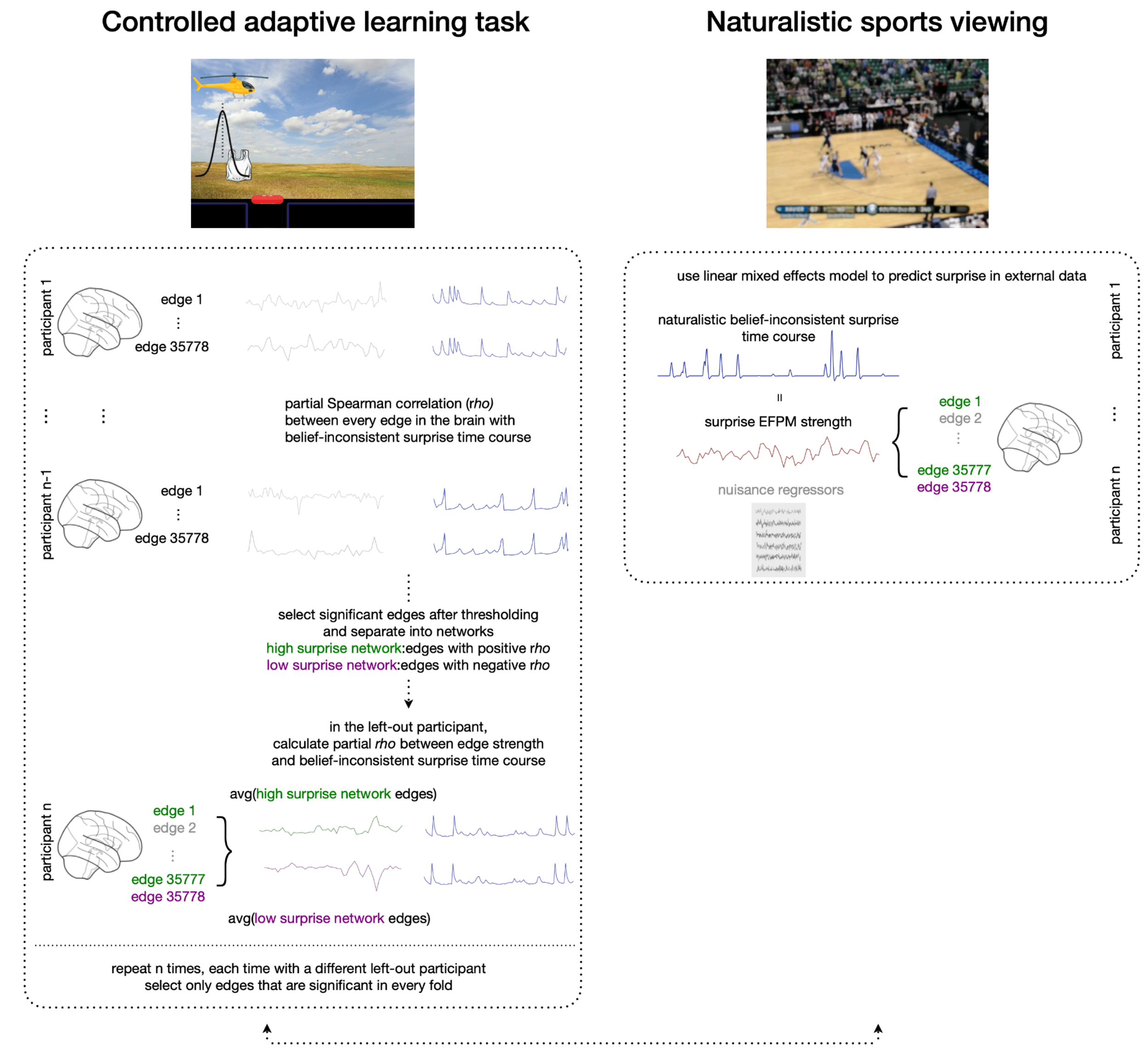


**Supp Fig 1.** EFPM-building methods overview. **Left**: An edge-fluctuation-based predictive model was trained to identify brain region pairs (edges) whose moment-to-moment co-activation varies with surprise in the controlled adaptive learning task. **Right**: Surprise EFPM generalizability was tested on the naturalistic sports viewing dataset. A mixed-effects model was used to predict belief-inconsistent surprise in the basketball video from surprise EFPM network strength and other regressors.

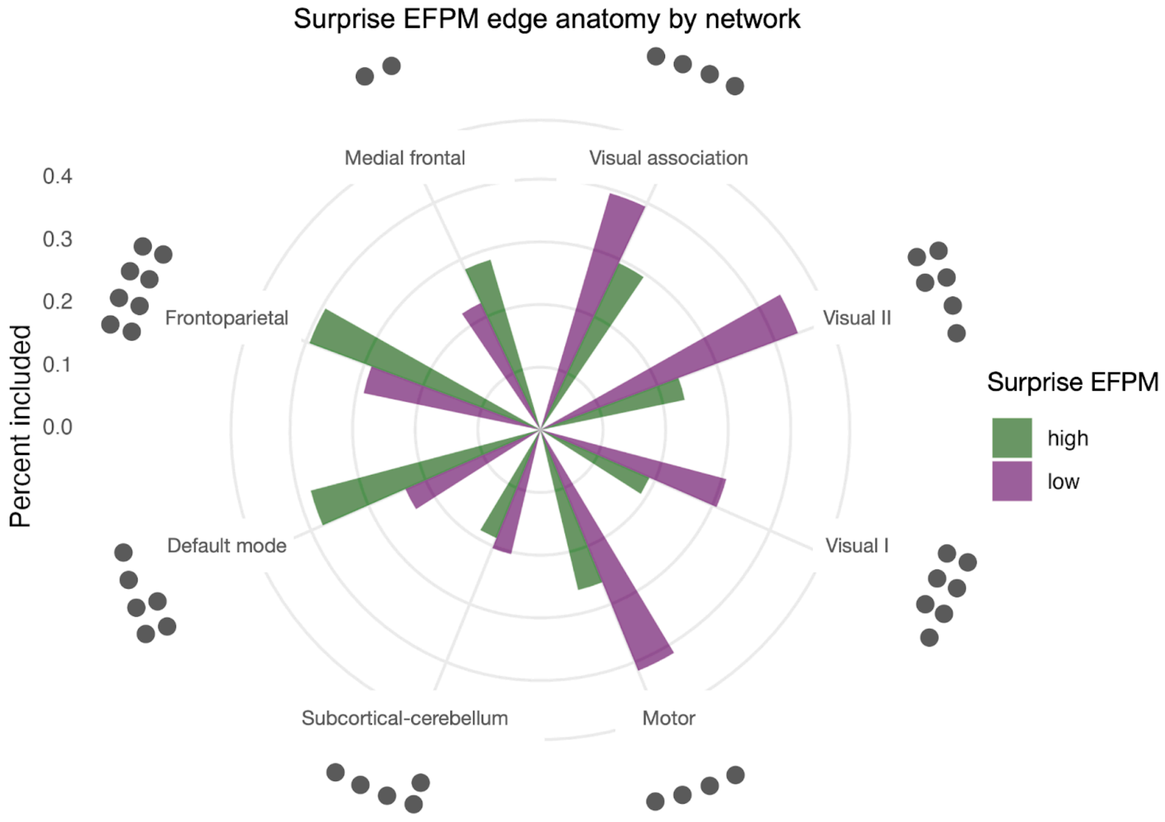


**Supp Fig 2.** Surprise EFPM anatomy defined by canonical resting-state networks. Number of dots indicates the importance of each network in the surprise EFPM anatomy. A dot was plotted each time the lesioning of edges within that network or between that network and another network led to failure of generalization to the external NCAA data.

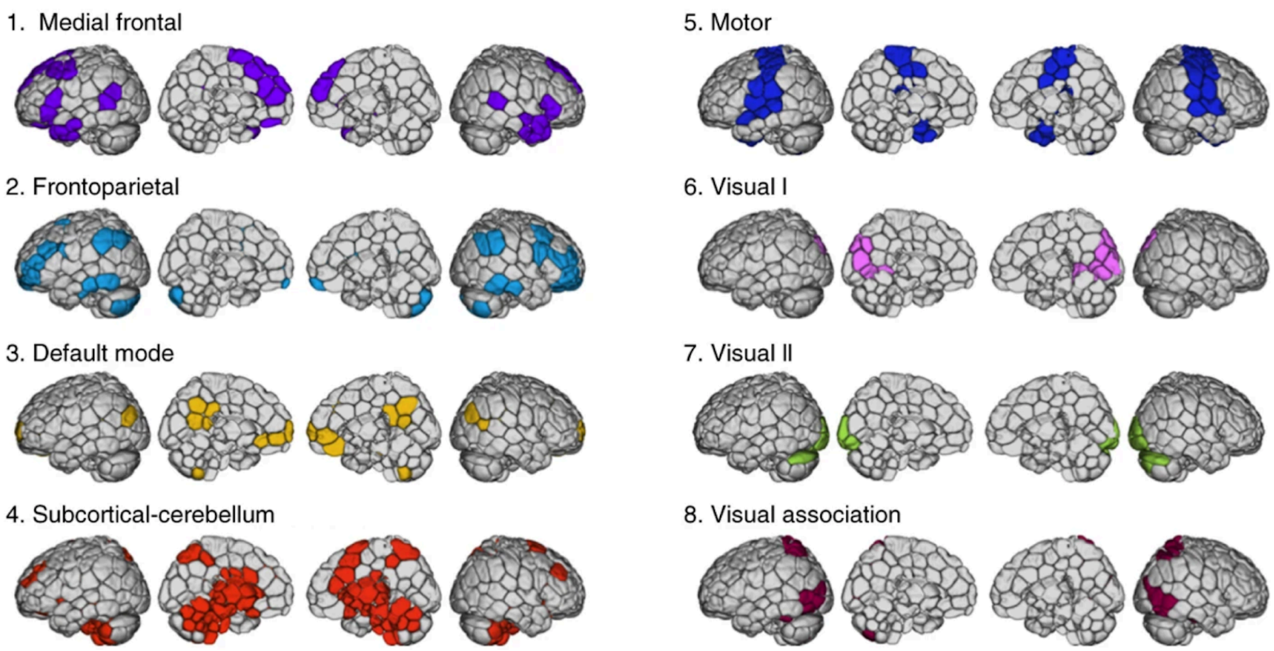


**Supp Fig 3.** Pre-defined functional networks from Finn et al. (2015). Figure adapted from Finn et al. (2015).

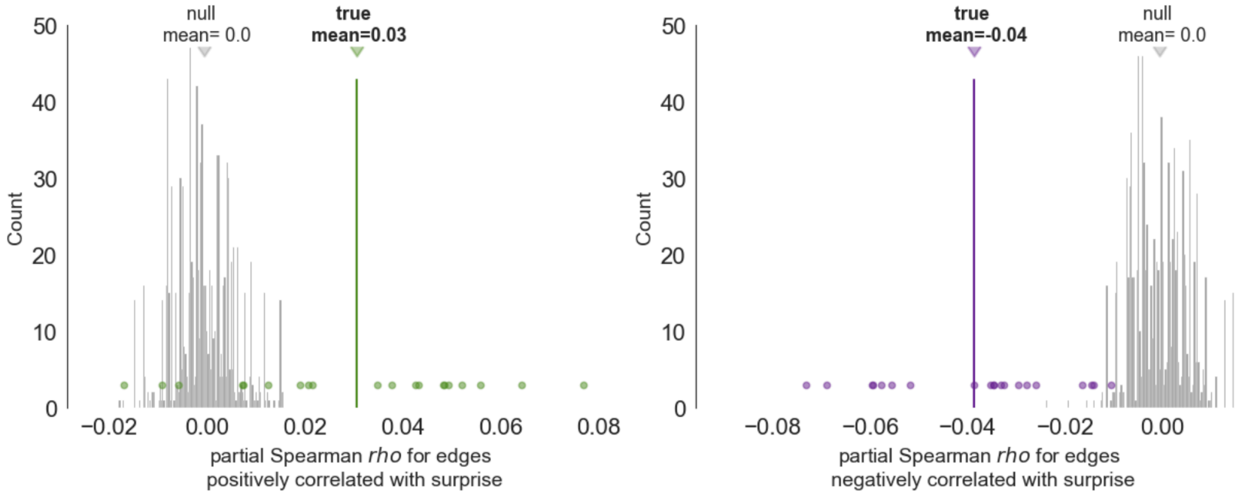


**Supp Fig 4.** An edge-fluctuation-based predictive model was also trained and validated in a flipped order: we trained on the naturalistic sports viewing dataset and tested for generalizability on the controlled adaptive learning dataset. Consistent with the main surprise EFPM, internal training significance was assessed by phase-randomizing the co-fluctuation strength time series for the positive and negative correlated edges in the NCAA training fold. Edge selection after all 20 LOOCV folds was consistent with the main surprise EFPM. The same mixed-effects model as specified for the adaptive learning task (*Comparing the generalizability of the surprise EFPM with that of the predefined networks* in Methods) was used, besides changing the brain network score term from that of the predefined networks to that of the network obtained from training on belief-inconsistent surprise in the NCAA videos. Significance was assessed with the same circular shifting approach described in *Methods*. **Left**: The observed partial Spearman *rho* value for edges positively (green distribution) correlated with belief-inconsistent surprise (mean within-subject partial *rho*=0.03; two-tailed *p*=1/1001); **Right**: or negatively (purple distribution) correlated with belief-inconsistent surprise (mean within-subject partial *rho*=-0.04; two-tailed *p*=1/1001). The null distributions (gray distributions) were obtained from repeating the internal CV 1000 times with phase-randomized summary edge score and then averaged across these repetitions. Importantly, flipping the order of training and testing datasets still preserved the generalizability of the model (*ß*=0.005*, t*(14862.32)=3.580, *p*=0.004), suggesting that the EFPM approach is successful in identifying neural correlates of belief-inconsistent surprise across contexts.

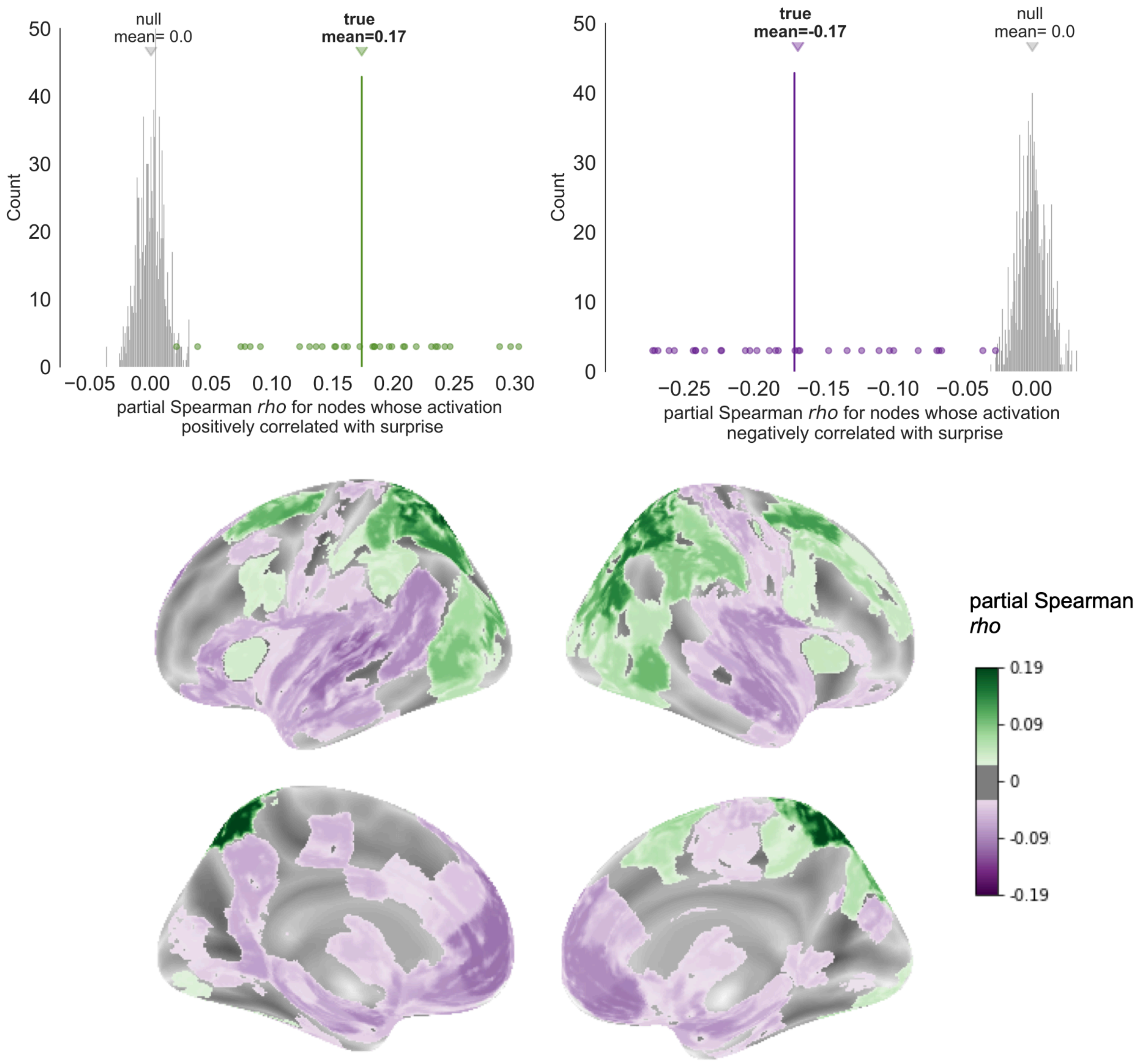


**Supp Fig 5.** Region-level BOLD activation. **Top**: Results from the internal cross-validation model training process on region level activations. The observed partial Spearman *rho* value for the high (left, green distribution) and low (right, purple distribution) surprise network was plotted with the null distribution (gray distribution) obtained from repeating the internal LOOCV 1000 times with phase-randomized regional activation score and then averaged across subjects. **Bottom**: Brain regions predicting belief-inconsistent surprise identified from regional BOLD activity.

*Number of edges in canonical networks*

| | medial frontal | frontoparietal | default mode | subcortical/ cerebellum | motor | visual I | visual II | visual association |
|---|---|---|---|---|---|---|---|---|
| medial frontal (29 nodes) | 406 | 986 | 580 | 2,610 | 1,450 | 522 | 261 | 522 |
| frontoparietal (34 nodes) | | 561 | 680 | 3,060 | 1,700 | 612 | 306 | 612 |
| default mode (20 nodes) | | | 190 | 1,800 | 1,000 | 360 | 180 | 360 |
| subcortical/cerebellum (90 nodes) | | | | 4,005 | 4,500 | 1,620 | 810 | 1,620 |
| motor (50 nodes) | | | | | 1,225 | 900 | 450 | 900 |
| visual I (18 nodes) | | | | | | 153 | 162 | 324 |
| visual II (9 nodes) | | | | | | | 36 | 162 |
| visual association (18 nodes) | | | | | | | | 153 |
| total edges in each network | 7337 | 8517 | 5150 | 20025 | 12125 | 4653 | 2367 | 4653 |

**Supp Table 1.** Number of edges that belong to each of the pre-defined networks in Finn et al. (2015). The sum of the upper triangular of the matrix is the total number of unique edges, 35,778.

*Surprise EFPM external testing results*

| variable | Estimate | t_value_observed | *p* | t_value_null_mean |
|---|---|---|---|---|
| **surprise EFPM strength** | **0.04** | **3.95** | **.047*** | **0.07** |
| game remaining | -0.12 | -12.09 | .371 | -0.21 |
| home possession | 0.06 | 6.45 | .363 | -0.61 |
| court position | -0.08 | -8.36 | .288 | -0.05 |
| **luminance** | **-0.47** | **-38.67** | **.001**** | **-0.55** |
| prosody | -0.03 | -2.95 | .769 | -0.65 |
| **video motion** | **0.30** | **29.27** | **.001**** | **-0.18** |
| auditory envelop | 0.11 | 5.79 | .436 | -3.30 |
| framewise displacement | 0.01 | 0.65 | .599 | 0.00 |

*Note.* All p values were two-tailed. * p < .05, ** p < .01, *** p < .001

**Supp table 2.** Results from the linear mixed-effects model predicting external belief-inconsistent surprise in the NCAA dataset.

*External testing model results excluding video luminance and video motion*

| variable | Estimate | t_value_observed | *p* | t_value_null_mean |
|---|---|---|---|---|
| **surprise CFPM strength** | **0.04** | **4.29** | **.032*** | **0.07** |
| **game remaining** | **-0.16** | **-16.89** | **.031*** | **-0.64** |
| home possession | 0.06 | 6.04 | .425 | -3.31 |
| court position | -0.05 | -5.24 | .716 | -0.31 |
| prosody | -0.06 | -5.42 | .498 | 0.02 |
| auditory envelop | 0.20 | 10.02 | .176 | -0.65 |
| framewise displacement | 0.01 | 0.47 | .696 | 0.00 |

*Note.* All p values were two-tailed. * $p < .05$, ** $p < .01$, *** $p < .001$

**Supp table 3.** Results from the linear mixed-effects model predicting external belief-inconsistent surprise in the NCAA dataset, excluding lower level visual regressors (video luminance and video motion).

*Dynamic functional connectivity model vs. surprise EFPM training results*

| window size | | within subject *rho* | *p* value | mean null *t* value | generalization *p* value |
|---|---|---|---|---|---|
| Surprise EFPM | pos | 0.09 | 0.001** | 0.00 | 0.047* |
| | neg | -0.10 | 0.001** | 0.00 | |
| 11 TRs | pos | 0.001 | 0.474 | -0.001 | 0.102 |
| | neg | 0.010 | 0.226 | -0.000 | |
| 21 TRs | pos | -0.012 | 0.768 | 0.001 | 0.091 |
| | neg | 0.009 | 0.286 | 0.000 | |
| 31 TRs | pos | -0.018 | 0.843 | 0.000 | 0.415 |
| | neg | 0.011 | 0.289 | 0.000 | |

*Note*. All *p* values were two-tailed. * $p < .05$, ** $p < .01$, *** $p < .001$. Training was performed on the controlled task dataset (left), and generalization results were obtained from the naturalistic video dataset (right). TR=2.5s in the controlled task dataset.

**Supp table 4.** Comparison of results from building models on edge co-fluctuations vs. sliding-window functional connectivity. Three window sizes were used here (11, 21, 31 TRs, corresponding to 27.5, 52.5, and 77.5 seconds). A model built from moment-to-moment edge fluctuations better-generalized both within and across datasets.

*Canonical network predicting belief-inconsistent surprise*

| Dataset | network name | estimate | *t* value | *p* value | mean null *t* value |
|---|---|---|---|---|---|
| Controlled adaptive learning task | saCPM | -0.01 | -3.29 | 0.01 | 0.00 |
| | MF_MF | -0.01 | -3.91 | 0.00 | 0.02 |
| | FP_FP | -0.00 | -2.41 | 0.04 | 0.02 |
| | FP_DMN | 0.01 | 3.30 | 0.00 | -0.01 |
| | FP_SubcortCere | 0.00 | 2.71 | 0.02 | -0.02 |
| | FP_VA | 0.00 | 2.23 | 0.04 | -0.01 |
| | DMN_DMN | -0.01 | -4.85 | 0.00 | 0.02 |
| | DMN_VA | 0.01 | 4.36 | 0.00 | -0.01 |
| | SubcortCere_SubcortCere | -0.00 | -3.06 | 0.01 | 0.03 |
| | SubcortCere_VI | -0.00 | -2.40 | 0.03 | 0.01 |
| | Motor_VA | -0.01 | -3.95 | 0.00 | 0.01 |

*Note.* All p values were two-tailed. * $p < .05$, ** $p < .01$, *** $p < .001$

**Supp table 5.** Canonical networks predicting belief-inconsistent surprise. Only significant networks are shown. Network abbreviations: saCPM: sustained attention connectome-based predictive model; MF: Medial frontal; FP: Frontoparietal; DMN: Default mode; SubcortCere: Subcortical-cerebellum; Motor: Motor; VI: Visual I; VII: Visual II; VA: Visual association.
